# The Hippo-YAP/TAZ-TEAD axis promotes multipotency in bone marrow stromal cells

**DOI:** 10.1101/2024.05.14.594149

**Authors:** Yu-Hao Cheng, Chun Hei Ryan Chan, Qin Bian, Randall K. Merling, Sergei A. Kuznetsov, Pamela G. Robey, Patrick Cahan

## Abstract

The mechanisms by which bone marrow stromal cells (BMSCs) maintain multilineage potency *in vitro* remain elusive. To identify the transcriptional regulatory circuits that contribute to BMSC multipotency, we performed paired single-nucleus multiomics of the expansion of freshly isolated BMSCs and of BMSCs undergoing tri-lineage differentiation. By computationally reconstructing the regulatory programs associated with initial stages of differentiation and early expansion, we identified the TEAD family of transcription factors, which is inhibited by Hippo signaling, as highly active in the BMSC *in vitro* multipotent state. Pharmacological inhibition of TEAD enhanced BMSC osteogenic and adipogenic differentiation, whereas its activation maintained BMSCs in an undifferentiated state, supporting a model whereby isolation of BMSCs coincides with a TEAD-controlled transcriptional state linked to multipotency. Our study highlights the Hippo pathway as a pivotal regulator of BMSC multipotency, and our regulatory network inferences are a reservoir of testable hypotheses that link transcription factors and their regulons to specific aspects of BMSC behavior.

## Introduction

Bone marrow stromal cells (BMSCs) are a diverse group of cells that play crucial roles in both bone homeostasis, by regulating the balance of osteoblast^1,2^ and osteoclast activity^3^, and in hematopoiesis, by providing and maintaining a supportive niche^4–8^. Both roles are dependent on the stem cell properties of BMSCs: their ability to self-renew and to produce functional, distinct cell types of the bone marrow stroma. Lineage tracing studies have provided *in vivo* evidence of BMSCs contributing to bone formation ^9^ and the maintenance of hematopoiesis ^10,11^. This evidence is further supported by the observed formation of ossicles and bone marrow following subcutaneous transplantation of freshly isolated BMSCs ^12,13^. The more accessible *in vitro* assays are frequently used to explore BMSC biology. *Ex vivo* plating of BMSCs in plastic dishes allows a subset of them to adhere to the vessel surface to form fibroblast colonies or CFU-Fs^14,15^. After clonal expansion in culture, BMSCs can be coaxed to differentiate into chondrocytes, osteoblasts, and adipocytes, and they can self-renew, as demonstrated by generation of ectopic bone marrow upon serial subcutaneous transplantation^5^.

The molecular mechanisms that underpin these properties have been a subject of intensive and long-standing investigation, not least because their dysregulation is a major contributor to pathologies of the skeletal and hematopoietic systems^16–18^. In particular, much has been learned about the signaling and transcriptional regulators of lineage differentiation of BMSCs *in vitro*. Specifically, mouse BMSCs can be differentiated *in vitro* to adipocytes using either a cocktail of insulin, dexamethasone, and isobutyl methylxanthine (IBMX)^19^ or by rosiglitazone^20^, which triggers up-regulation of PPARγ and C/EBPα, which in turn up-regulate expression of adiponectin, and FABP4^21,22^. Chondrogenic differentiation can be induced by transforming growth factor-beta (TGF-β)^23,24^, while osteogenic differentiation can be triggered by bone morphogenetic proteins (BMPs)^25^, vitamin D3^26^, and ascorbic acid^27,28,29^.

Less is known about the identity and role of signaling pathways and transcriptional circuits in establishing and maintaining multipotency. Several genes and proteins have been implicated in the multipotent state based on correlative observations. For example, Sox11 expression in undifferentiated, cultured BMSCs correlates with bone-forming potential upon differentiation^30^. Similarly, the extracellular proteins *Loxl2*, *Tgfbi*, and *Serpine2* are higher expressed in undifferentiated BMSCs than in differentiated progeny^31^, suggesting that they contribute to multipotency or self-renewal. How these molecules are linked to multipotency remains unclear, and more broadly, we are aware of no mechanistic model that adequately explains BMSC multipotency.

Here, we set out to identify transcriptional contributors to BMSC multipotency by single-nucleus multiomics (snMultiomics). We analyzed BMSCs during their tri-lineage differentiation and early expansion phases. The analysis reaffirmed the roles of well-known regulators such as *Pparg* and *Runx2* in BMSC differentiation. More importantly, we identified TEAD as critical factor for the expansion and maintenance of BMSC multipotency. This discovery underscores the crucial role of Hippo pathway in BMSC multipotency and provides a basis for developing more targeted strategies to control BMSCs.

## Results

### Multiomics Profile of Tri-lineage Differentiating Multipotent BMSCs

BMSCs exhibit tri-lineage multipotency, a characteristic extensively studied *in vitro*. However, accurately identifying the multipotent BMSCs *in vivo* and *in vitro* has been a subject of debate^32,33^. Direct evidence depicting the differentiation trajectories would enhance our understanding of the characteristics of the sourcing multipotent populations^34,35^. To address this question, we conducted a multi-seq multiplexed snMultiomics experiment on the early differentiation BMSCs *in vitro*. Simultaneous capture of both ATAC-seq and RNA-seq at the single-nucleus level allowed us to precisely identify not only the active genes but also the active chromatin regions where transcription factors (TFs) bind. Such insights were crucial for decoding the transcriptional networks driving multipotent BMSC differentiation and sustaining their multipotency.

The BMSC early differentiation experiment involved sampling multiple timepoints throughout the process for each lineage, starting from the initiation to the seventh day of differentiation. For each lineage, samples of different timepoints were prepared days in advance to ensure that samples belonging to the same lineage could be collected simultaneously (Figure 1A). This approach aimed to minimize the potential influences resulting from the subsequent nuclei isolation, transposition, and single-nucleus preparation processes. After excluding likely doublets and low-quality nuclei, we captured a total of 8169 nuclei from three lineages, which each lineage contributing about one third of the total number (Figure S1A). To visualize the multiomics data more comprehensively, we calculated the joint embedding by integrating both the transcriptomic and the chromatin accessibility information. After batch correction using the Harmony algorithm, the nuclei organized into three distinct transcriptomic clusters (Figure S1B), which could be further subdivided into eight transcriptomic subclusters (Figure 1B). Additionally, these cells display three distinct epigenetic clusters (Figure 1C).

**Figure 1:**
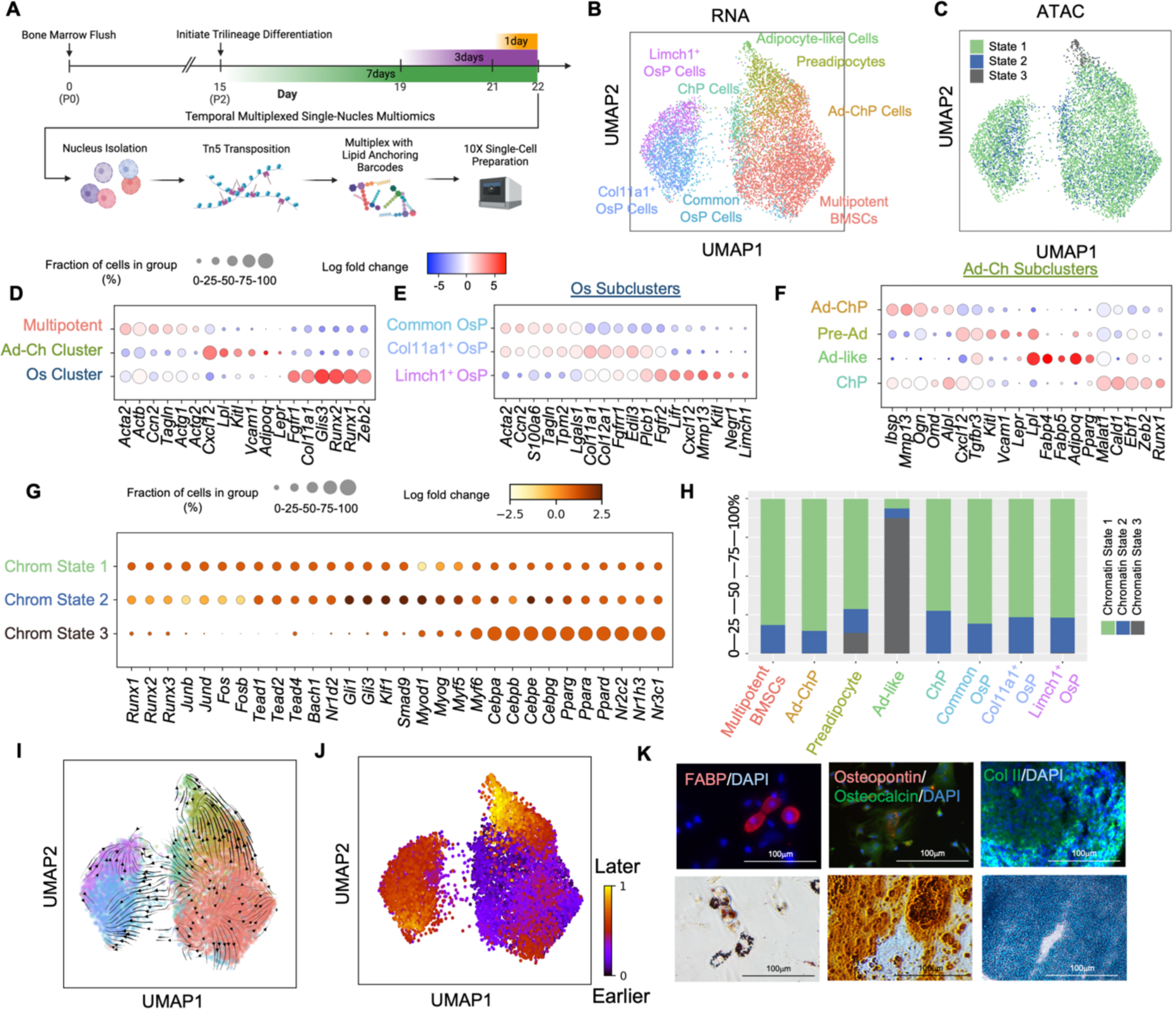
Multiomics Profiles of Early Differentiating Multipotent BMSCs. (A) Schematic of the multiplexed snMultiomics design. (B) Eight transcriptomic distinct populations were discovered, each belonging to either the adipogenic, chondrogenic, or osteogenic lineage. (C) Three epigenetic distinct chromatin states were uncovered. (D) The differentially expressed genes in three major transcriptomic distinct cluster. (E) Identification of genes differentially expressed in osteogenic subclusters. (F) Identification of genes differentially expressed in adipochondrogenic subclusters. (G) The top enriched motifs in each distinct chromatin state. (H) The attribution of four different chromatin states in each transcriptomic distinct cluster. (I) RNA velocity analysis revealed cell trajectories, guiding the commitment of multipotent BMSCs toward downstream endpoints. (J) The latent time, derived from the RNA velocity analysis. (K) Assays verified the multipotent ability in multipotent BMSCs to differentiate into functional adipocyte, osteoblast, and chondrocyte-like cells *in vitro*.

At the transcriptional level, the largest cluster (*n* = 3763) was marked by high expression of cytoskeletal-associated genes *Tagln*, *Acta2*, *Actb*, *Actg1*, *Actg2*, and the connective tissue growth factor *Ccn2* (Figure 1D). This cluster comprised 86% of the nuclei from day 0, which represented a common origin for all three lineages. Therefore, we designated it as “multipotent BMSCs” (Figure S1B). The second cluster showed enrichment in *Cxcl12* and several adipogenic markers, including, *Lpl*, *Lepr*, and *Adipoq* (Figure 1D). Predominantly present in adipogenic and chondrogenic lineages, it was termed the “adipo-chondrogenic cluster” (Figure S1B). The last cluster demonstrated enrichment in many early osteogenic markers, like *Fgfr1*, *Col11a1*, and *Runx2* (Figure 1D), and was hence termed the “osteogenic cluster” (Figure S1B).

The osteogenic cluster further divided into three subclusters. One of these subclusters showed enrichment in collagens (*Col11a1*, *Col12a1*) and FGF receptors (*Fgfr1*, *Fgfr2*), and we have identified it as the “Col11a1^+^ osteoprogenitor cells” (Figure 1E). Another subcluster displayed enrichment in Limch1 and certain CAR cell markers, including *Cxcl12* and *Kitl*, and has been designated as the “Limch1^+^ osteoprogenitor cells” (Figure 1E). The remaining subcluster, positioned between the multipotent BMSCs and the two osteoprogenitor clusters, exhibited fibroblastic markers, including *Tagln*, *Timp1*, *S100a6*, and *S100a11*, and we have labeled it as the “common osteoprogenitor cells” (Figure 1E).

The adipo-chondrogenic cluster further divided into four subclusters. One of these subclusters displayed notable enrichment in adipocyte-related markers, such as *Fabp4*, *Fabp5*, *Adipoq*, *Adipoqr2*, *Lpl*, and *Lepr*, and we have labeled it as “adipocyte-like cells” (Figure 1F). Another subcluster was exclusive to the chondrogenic lineage and showing enrichment in early chondrogenic TFs, including *Ebf1*, *Zeb2*, and *Runx1*; designated as the “chondroprogenitor cells” (Figure 1F). The third subcluster located upstream of adipocyte-like cells and expressed markers like *Cxcl12*, *Kitl*, and *Lepr*, but lacking mature adipocyte markers; has been termed “pre-adipocytes”. The final subcluster displayed osteogenic markers, such as *Ibsp*, *Ogn*, and *Omd*, but also displayed lower expression of markers belonging to both the pre-adipocytes and the chondroprogenitor cells, such as *Cxcl12*, *Tgfbr3*, *Malat1*, and *Cald1*, has been named “adipo-chondrogenic progenitor cells” (Figure 1F).

Upon comparing the diverse transcriptomic subclusters identified during BMSC tri-lineage differentiation with the results of unsupervised clustering using ATAC information, we observed the presence of only three distinct epigenetic clusters, denoted as chromatin states 1-3 (Figure 1G). Specifically, Chromatin state 1 was predominantly marked by motifs associated with AP-1 factors, such as *Fos* and *Fosb*, as well as *Runx* factors. Chromatin state 2 was characterized by a notable presence of motifs related to *Tead* and *Bach*, along with *Gli* factors. In contrast, chromatin state 3 displayed significant enrichment in *Cebp* and *Ppar* factors. Further analysis of these chromatin states across the different transcriptomic clusters showed that adipocyte-like cells primarily aligned with chromatin state 3. Meanwhile, the other clusters demonstrated a mix of chromatin states 1 and 2, suggesting a dual epigenetic landscape within these cell types (Figure 1C).

While comparing the clusters observed from the initial day to the endpoints in each lineage, we were able to infer the cell identity and global trajectory from the undifferentiated multipotent BMSCs via intermediate states toward the termination (Figure S1C). All three lineages shared the same origin as the multipotent BMSCs, and soon the cells bifurcated into two distinct trajectories. One went toward adipo-chondrogenic progenitor cells, serving as the upstream precursor of both adipogenic and chondrogenic lineages, and the other entered a state of common osteoprogenitor cells (Figure 1B, S1A-C). While comparing the attribution of clusters in each timepoint, we were able to evaluate the relative speed of multipotent BMSCs committing to different lineages. On day one, the adipogenic differentiation had already shown a substantial proportion committed toward either adipo-chondrogenic progenitor cells or preadipocytes (Figure S1D). The osteogenic lineage didn’t show a transition toward common osteoprogenitor cells until day three (Figure S1E). The chondrogenic differentiation stayed at the adipo-chondrogenic progenitor cells even on day seven (Figure S1F). The RNAvelocity depicted a consistent flow originating from the multipotent BMSCs toward endpoints of three different lineages (Figure 1I), and the latent time analysis showed consistent results compared to the true timepoints we captured for each sample (Figure 1J).

To assess the potential of precursor cells to develop into mature cells *in vitro* with similar function as their *in vivo* counterparts, we conducted functional staining on the fully differentiated cells. The adipocyte-like cells exhibited enhanced expression of FABP and intracellular lipid accumulation, demonstrated with Oil red O staining. In contrast, the osteogenic cells displayed positive expression of osteopontin (OCN/SPP1) and osteocalcin (BGLAP), accompanied by the deposition of calcium, which was visualized as red staining in Alizarin Red staining. During chondrogenic differentiation, chondrocyte-like cells were generated, expressing type II collagen and displaying acidic mucopolysaccharides, as indicated by Alcian blue staining. These findings strongly suggest that the precursor cells we observed exhibit a certain degree of functional similarity to their *in vivo* counterparts (Figure 1K).

### Regulatory Programs Governing Fate Decisions of Multipotent BMSCs

From the BMSC early differentiation experiment, we successfully characterized distinct multiomic states of multipotent BMSCs *in vitro* and identified intermediate transcriptomic states along the lineage paths. This prompted us to explore the molecular signatures enriched in each state to decipher the regulatory programs that drive lineage specification and that sustain multipotency. To accomplish this, we performed network analyses to score the activity of TFs and link them to their predicted targets relevant to each transcriptomic state or lineage. Our initial step involved evaluating the motif activity in each transcriptomic state, which allowed us to identify TFs actively engaging with their downstream target genes, thereby refining our search to accurately identify these crucial regulators.

Multipotent BMSCs were found to be highly enriched in AP-1 motifs, including *Fosb*, *Fos*, *Jund*, *Junb*, along with other distinct motifs such as *Tead1*, *Tead2*, *Runx3*, and *Bach2*. The TFs exhibiting high motif activity were likely to be actively involved in regulating fundamental properties of multipotent BMSCs, including cell migration, proliferation, and the maintenance of multipotency. In addition to these highly expressed and highly active factors, there were several other factors that exhibit high expression levels but did not demonstrate significant motif activity. For example, *Runx1*, *Runx2*, *Mef2a*, *Smad3*, *Vdr*, and *Zeb2* (Figure 2A, orange panel). The discrepancy between motif activity and expression level suggested that these factors might have been priming multipotent BMSCs to respond to external stimuli that had the potential to drive the cells toward differentiation.

**Figure 2:**
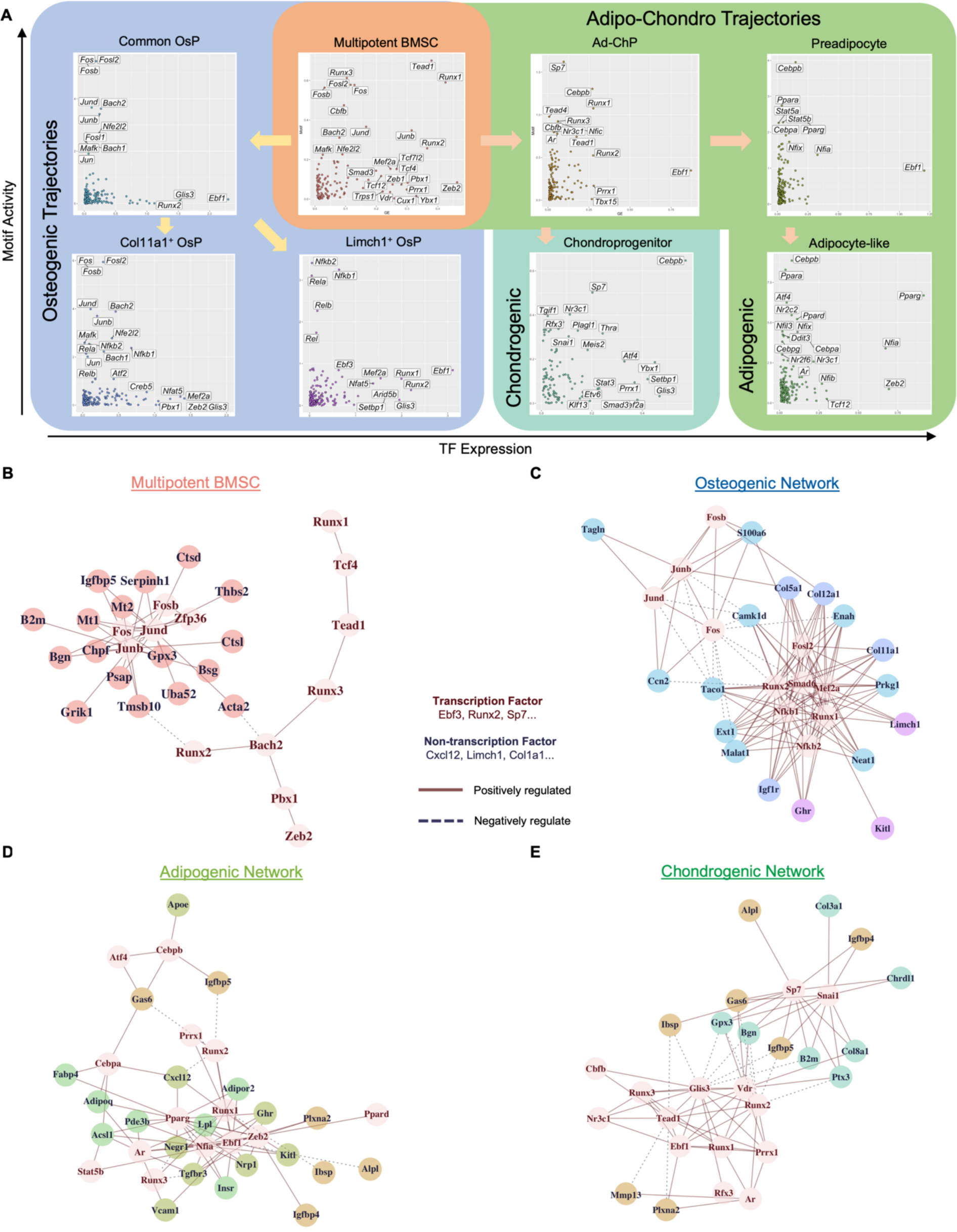
Gene Regulatory Networks Governing Multipotent BMSC Fate Decisions. (A) Simultaneous evaluation of expression and their motif activity of TFs enables the identification of critical regulators governing differentiating BMSC states. (B) Regulatory networks governing multipotent BMSCs. (C) Regulatory networks governing osteogenic fate commitment. (D) Regulatory networks governing adipogenic fate commitment. (E) Regulatory networks governing chondrogenic fate commitment.

In the osteogenic lineage differentiation, AP-1 factors, including *Fos*, *Jun*, *Junb*, were notably active in the common osteoprogenitor cells, although their expression levels were not as high as *Ebf1* or *Glis3*. The subsequent bifurcation of the osteogenic lineage was primarily driven by the upregulation of Nfkb-related factors and reduction of AP-1 activity. Specifically, the Nfkb-related factors, *Nfkb1*, *Nfkb2*, *Rela*, and *Relb,* were highly active in both Limch1^+^ and Col11a1^+^ osteoprogenitor cells; however, there was a significant suppression of AP-1 activity in Limch1^+^ osteoprogenitor cells. Additionally, apart from the most active motifs, we observed an increase in expression levels of *Runx1*, *Runx2*, *Mef2a*, and *Pbx1* in Col11a1^+^ osteoprogenitor cells, while *Ebf1* was upregulated in the Limch1^+^ osteoprogenitor cells (Figure 2A, blue panel). In the context of adipogenic lineage differentiation, specific motifs became more accessible during the transition to the adipo-chondrogenic progenitor cells, including *Sp7*, *Runx1*, and *Runx3*.

Simultaneously, there was an upregulation in the gene expression of *Cebpb* along with an increase in accessible motifs in the downstream adipogenic pathway. Besides *Cebpb*, motifs of *Ppara* and *Pparg* gradually increased throughout the trajectory of maturation of preadipocytes to adipocyte-like cells (Figure 2A, green panel). In the chondrogenic pathway, there was an increase in motifs of *Ppara*, *Pparg*, and *Ebf*, while there was a simultaneous upregulation in *Mef2a* and *Glis3* (Figure 2A, light green panel).

To uncover the regulatory programs governing each cell state and dynamic regulatory program along each lineage, we utilized “Epoch”, considered only transcriptomic information but captured temporal regulatory programs^36^, enabling the reconstructing dynamic networks for the adipogenic, chondrogenic, and osteogenic lineages.

In the multipotent BMSCs, our findings revealed a network with two prominent hubs: one centered around AP-1 factors and the other encompassing Runx factors, *Bach2*, and *Tead1* (Figure 2B). Among the diverse regulatory interactions, *Acta2* emerged as a noteworthy multipotent BMSC marker to highlight. *Acta2* was simultaneously regulated by both the AP-1 and *Bach2* hubs. Although there was no direct evidence confirming this regulatory relationship specifically in multipotent BMSCs, evidence from other cell types suggested a similar mechanism. For instance, *Acta2* expression had been reported to be suppressed during retrovirus-induced *Bach2* expression in multipotent hematopoietic progenitors^37^, and the AP-1 factor JUN was known to upregulate expression of *Acta2* in smooth muscle cells^38^.

For the osteogenic lineage, our dynamical network revealed that AP-1 factors were the center of the common osteoprogenitor cells, while the bifurcation of two downstream osteoprogenitor cells was mainly governed by *Runx1*, *Runx2*, *Nfkb*-related factors, and *Pparg* (Figure 2C). The dynamic network highlighted several regulatory interactions between the active TF and markers. Notably, interactions between *Runx2* and markers enriched in common osteoprogenitor cells, including *Malat1*, *Ext1*, and *Enah*, were found to be upregulated by *Runx2* in MC3T3 preosteoblast using Chip-seq data ^39^. While some of these interactions had been explored in BMSC related cell line, others had also been validated in other stem or progenitor cell types. For instance, *Limch1* and *Col11a2* were among the genes regulated by *Mef2a* control in the neural progenitor cells ^40^. *Col5a1* was shown to be downregulated when AP-1 activity was inhibited during the hemogenic endothelium differentiation of mouse embryonic stem cell ^41^.

In the adipogenic lineage, *Sp7*, *Cebpb*, *Atf4*, and *Nfic* were the TFs that correlated to the markers of adipo-chondrogenic progenitors, and subsequently shifted to adipogenic networks centered with *Pparg*, *Cebpa*, and *Nfia* (Figure 2D). Several adipocyte markers, including *Fabp4*, *Adipoq*, and *Acsl1,* were simultaneously regulated by *Pparg* and *Cebpa*, it was known that both factors orchestrating adipocyte biology, and *Fabp4*, *Adipoq*, *Acsl1* had been verified as upregulated targets by *Cebpa* Chip-seq ^42^ and *Pparg* Chip-seq ^43^ in 3T3-L1 preadipocyte.

The chondrogenic network involved *Prrx1*, *Runx1*, *Runx2*, *Runx3*, *Snai1*, and *Vdr* (Figure 2E). A study found that Smad4 deficiency impaired chondrocyte differentiation and prevented chondrocyte hypertrophy. When Smad4 was deficient, *Runx2*, *Runx3*, and *Sp7* were significantly down-regulated, suggesting these factors may have played a critical role in regulating chondrogenesis^44^. Beyond these TFs, the downstream targets surprisingly showed multiple markers related to osteoblast. Although type III collagen is related to chondrocytes, many of the downstream targets were more closely linked to matrixes modulation in osteogenesis, such as *Bgn*, *Alpl*, and *Ibsp*^45^.

### Distinct Multiomics Profiles of Freshly Isolated BMSCs

Next, we explored the extent to which the culture and expansion of freshly isolated BMSCs was controlled by similar regulatory programs as those that we observed in the in vitro multipotent state (Figure 3A). To enrich for BMSC populations, we utilized fluorescence-activated cell sorting (FACS) to exclude hematopoietic and endothelial cells by targeting specific cell surface markers, including TER119, CD45, and PECAM1 (Figure S2A). We implemented stringent quality control measures for gene expression and epigenetic signatures, resulting in the inclusion of 8,634 high-quality nuclei from day 0, day 1, and day 3 post-isolation samples. To assess the consistency of BMSCs across different preparation batches, we employed multiplexing, tagging the post-isolation day 3 sample with day 0 and day 1 samples. Our analysis revealed that not only did the multiplexed samples correctly cluster with the independently submitted day 0 and day 1 samples in the UMAP embedding, but supervised classification using SingleCellNet also confirmed the assignment of time points. This suggests that technical variations between experiments are minimal (Figure S2B and Figure S2C).

**Figure 3:**
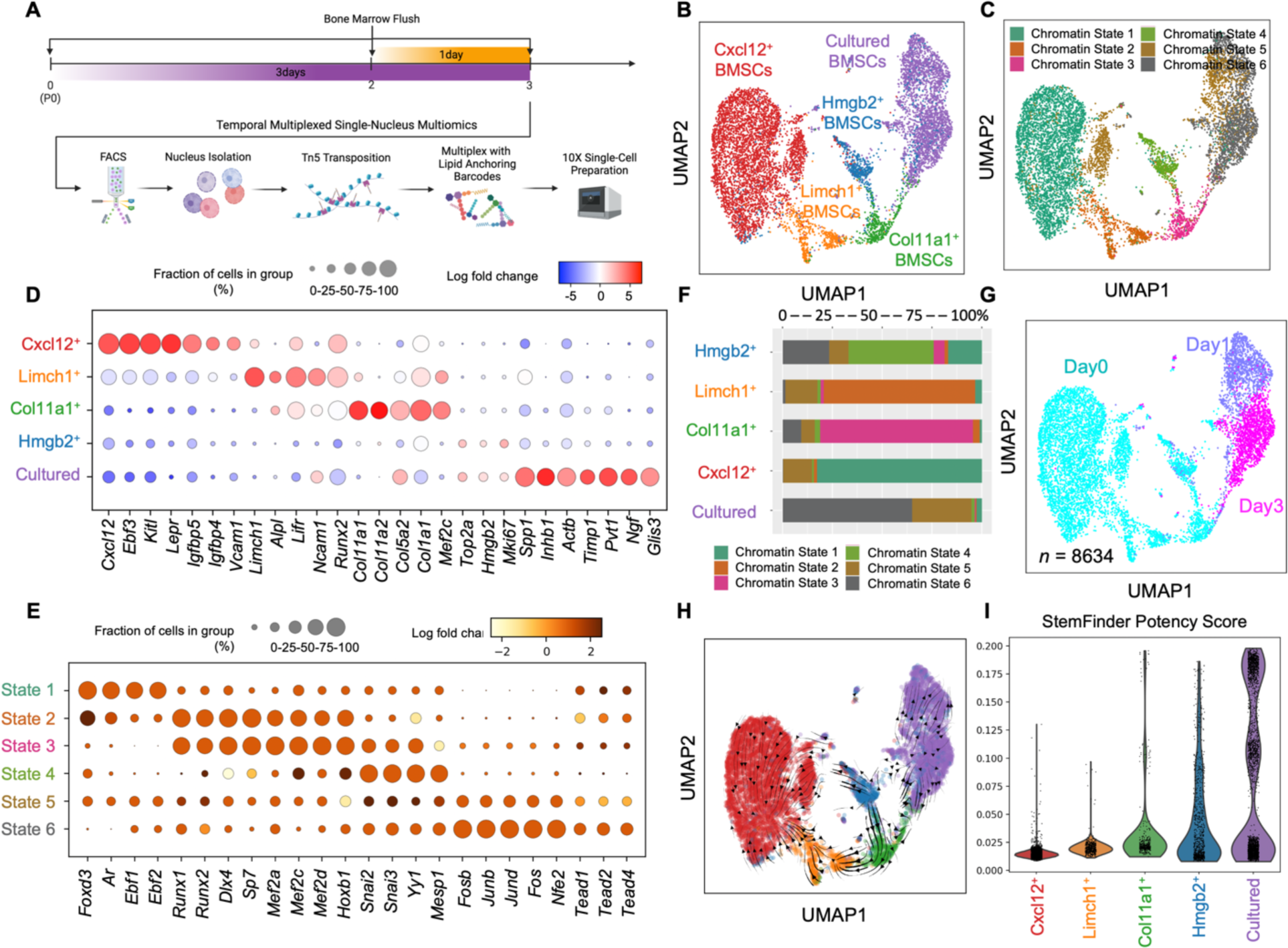
Multiomics Profile of BMSCs *in vivo* and *in vitro*. (A) The snMultiomics design was employed to capture BMSCs *in vivo* as well as early expansion. (B) Five transcriptomic distinct BMSC clusters were identified. (C) Identification of six epigenetic distinct chromatin states in BMSCs. (D) Differentially expressed genes enriched in each distinct cluster. (E) Each chromatin state displayed distinct differential motif activities. (F) The assignment of chromatin states to each transcriptomically distinct BMSC. (G) Time labels revealed rapid condensation of heterogenous BMSCs post-isolation. (H) RNA velocity analysis illustrates the flow of cells, indicating that Hmgb2^+^ BMSCs are a critical source of origin. (I) StemFinder predicts Hmgb2^+^ BMSCs as the most potent cells among other clusters *in vivo*, while cultured BMSCs *in vitro* exhibit a similar potency.

After conducting unsupervised clustering, we identified five distinct clusters at the transcriptomic level (Figure 3B) and six distinct clusters at the epigenetic level (Figure 3C). In terms of transcriptomic profiles, the Cxcl12^+^ BMSCs cluster expressed genes associated with Cxcl12-abundant reticular cells, including *Cxcl12*, *Ebf3*, *Lefr*, the hematopoietic support factor *Kitl,* and adipogenic marker *Igfbp5*. The Limch1^+^ BMSCs cluster expressed genes such as *Limch1*, the mineralization enzyme *Alpl* and the osteogenic master regulator *Runx2*. The Col11a1^+^ BMSCs expressed multiple types of collagen crucial for building the backbone of extracellular matrix, including type I, type V, and type XI. The Hmgb2^+^ BMSCs cluster showed enrichment of *Hmgb2* and other genes related to the cell cycle, including *Top2a* and *Mki67*. The final cluster displayed a few genes associated with Spp1^+^ BMSCs, such as *Spp1* and *Timp1*, but also expressed several distinct genes, including *Inhb1*, *Actb*, *Pvt1*, *Ngf*, and *Glis3* (Figure 3C and Figure 3D). Notably, this cluster was predominantly observed on days 1 and 3 post-isolation, which we termed as the "cultured BMSCs" (Figure S2D).

At the epigenetic level, the first chromatin state was enriched for motifs such as *Ebf1*, *Ebf2*, *Ar*, and *Foxd3* (Figure 3E). The second and third chromatin states exhibited motifs associated with osteogenic regulators, including *Runx* factors, *Sp7*, and *Mef2* factors. The fourth chromatin state displayed enrichment of *Snai* factor motifs and *Mesp1*. The fifth chromatin state was enriched in AP-1 factors, encompassing *Fosb*, *Junb*, *Jund*, and *Fos*. The last chromatin state showed enrichment of *Nfe2* and *Tead* factors. Upon comparing the transcriptomic clusters with their underlying chromatin states, it became evident that each cluster corresponded to a specific chromatin state. For instance, Cxcl12^+^ BMSCs predominantly comprised chromatin state 1, Col11a1^+^ BMSCs had a higher fraction of chromatin state 3, Limch1^+^ BMSCs were associated with chromatin state 2, Hmgb2^+^ BMSCs exhibited a higher fraction of chromatin state 4, and cultured BMSCs were highly enriched in chromatin state 6. The only exception was chromatin state 5, which was distributed across all clusters but occupied a small fraction in each cluster (Figure 3F).

When the temporal information was considered, it became clear that the samples from day 1 and day 3 predominantly consist of a single BMSC cluster, whereas the day 0 duplicates exhibited multiple distinct BMSC clusters (Figure 3G). These findings suggested a rapid loss of BMSC heterogeneity immediately following isolation, with this transition occurring within a 24-hour timeframe. To assess the origin of the cultured BMSC population, we conducted trajectory analysis using RNA velocity. The results demonstrated that Hmgb2^+^ BMSCs not only exhibited a tendency to differentiate into Cxcl12^+^ BMSCs, Limch1^+^ BMSCs, and Col11a1^+^ BMSCs, but they also displayed a strong flow towards the cultured BMSC population (Figure 3H). This suggested that Hmgb2^+^ BMSCs were the likely source of the cultured BMSCs. Subsequently, we utilized StemFinder to address the inconsistency in predicting the most potent BMSC population between the meta-analysis and *in vitro* differentiation experiment. The results showed that Hmgb2^+^ BMSCs are predicted to be the most potent population within freshly isolated clusters, consistent with the findings from the meta-analysis. Intriguingly, the cultured BMSCs were found to have potency levels that are comparable to, or potentially even greater than, those of Hmgb2^+^ BMSCs (Figure 3I and Figure S2E). Both Hmgb2^+^ BMSCs and cultured BMSCs exhibited substantial features indicative of the S/G2M cell cycle phase, suggesting that both populations are actively dividing (Figure S2F).

To evaluate the capacity of cultured BMSCs to develop mature, terminally differentiated cells with similar functional features as their *in vivo* counterparts, we conducted *in vivo* transplantation. After a 10-week incubation period, we harvested the samples for functional assays. The transplants exhibited radiopaque ossicles (Figure S3A), and the HE staining revealed bone marrow-like structure in the middle (Figure S3B). FABP staining (Figure S3C) and Oil Red O (Figure S3D) confirmed the presence of mature adipocytes, while osteopontin (OCN/SPP1) and osteocalcin (BGLAP) staining (Figure S3E), along with calcium deposition in Alizarin Red staining (Figure S3F), confirmed the presence of active osteoblasts. Additionally, positive staining for type 2 Collagen (Figure S3G) and Alcian Blue staining indicating acidic mucopolysaccharides confirmed the presence of chondrocytes (Figure S3H). Y chromosome painting substantiated that the cells in the transplant originated from the male donor mice and not from the female recipients (Figure S3I). These findings strongly suggested that the cultured BMSCs collected on Day 3 were fully potent in deriving all three lineages *in vivo*.

### Multiomics Pinpoints Critical Regulators of BMSCs *in vivo* and *in vitro*

The early expansion of BMSCs revealed a rapid transition from the diverse freshly isolated BMSCs to a more homogenous multipotent BMSCs *in vitro*. This transition was marked by substantial changes in both gene expression and chromatin states as the cells adapted to the *in vitro* environment. This observation prompted us to explore what regulatory elements were activated in the cultured BMSCs as compared to those from freshly isolated BMSCs. To achieve this goal, we assessed the TF activity in the early expansion BMSCs and reconstructed the regulatory networks governing the cell state for each BMSC cluster.

To identify active TFs, we assessed the motif activity of expressed TFs within the five BMSC clusters. Cxcl12^+^ BMSCs displayed a higher activity of the *Ebf* factors and Ar (Figure 4A). Limch1^+^ and Col11a1^+^ BMSCs showed active TFs, including *Sp7*, *Runx1*, *Runx2*, *Mef2a*, and *Mef2c* (Figure 4A). Hmgb2^+^ BMSCs were primarily regulated by *Tal1*, and the cultured BMSCs, like the multipotent BMSCs characterized *in vitro*, showed significant activation of AP-1 TFs (Figure 2A and 4A). To map out the regulatory networks governing freshly isolated and early expanded BMSCs, we employed the "Pando" tool^46^. This method allowed us to consider both the motif activity and the differential gene expression, facilitating the reconstruction of a comprehensive regulatory network for each BMSC cluster.

**Figure 4:**
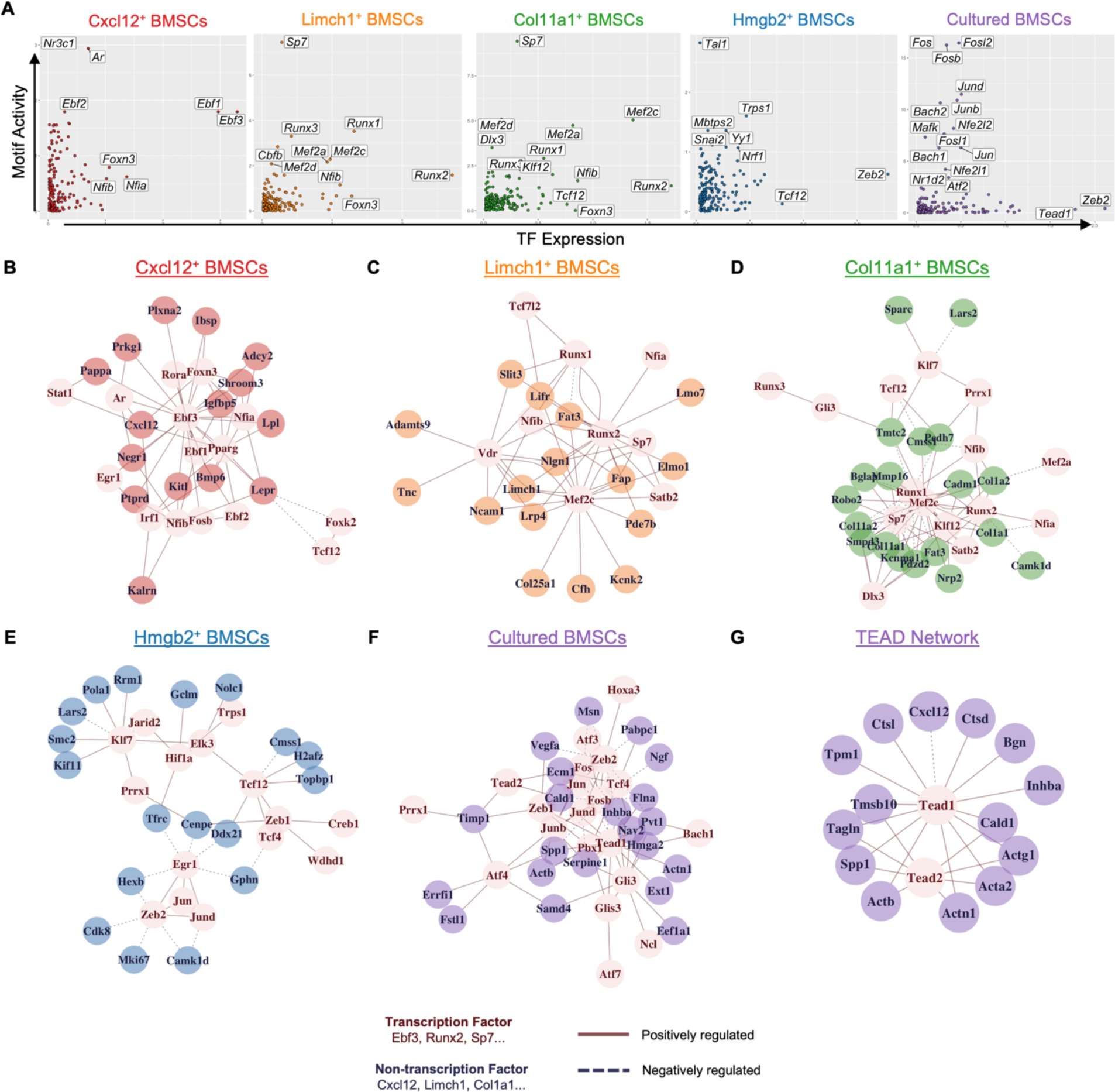
**Gene Regulatory Networks Governing BMSCs *in vivo* and *in vitro*** (A) Simultaneous assessment of expression and motif activity of TFs facilitates the identification of crucial regulators in BMSCs. (B) The gene regulatory networks that control Cxcl12^+^ BMSCs reveal significant regulations primarily involving *Ebf3* and *Pparg*. (C) The gene regulatory networks governing Limch1^+^ BMSCs are centered around *Runx2*, *Vdr*, and *Mef2c* as key regulators. (D) The gene regulatory networks governing Col11a1^+^ BMSCs comprise a central hub of *Mef2c*, *Runx1*, *Runx2*, and *Sp7*. (E) The gene regulatory networks governing Hmgb2^+^ BMSCs exhibit a loosely connected network involving AP-1 factors, *Tcf12*, and *Klf7*. (F) The gene regulatory networks governing cultured BMSCs primarily revolve around AP-1 factors and *Tead1* as central regulators. (G) The TEAD1 regulatory network governs multiple aspects related to actin filament formation.

In the Cxcl12^+^ BMSCs, *Ebf1* and *Ebf3* formed an interconnected hub that regulate multiple differentially expressed genes, including *Lepr*, *Kitl*, and *Igfbp5*. *Cxcl12*, on the other hand, was regulated by *Ar*, *Irf1*, *Pparg* and *Foxn3* (Figure 4B). Furthermore, *Mef2c*, *Runx1*, *Runx2*, and *Sp7* emerged as central hubs in the regulation of osteogenic BMSCs (Figure 4C and Figure 4D). However, the targets regulated by these factors differed between Limch1^+^ BMSCs and Col11a1^+^ BMSCs. For instance, *Runx2* directly regulated *Limch1* and *Lifr* in Limch1^+^ BMSCs, while *Runx2* directly regulated type I collagens and cell adhesion molecule *Cadm1* in Col11a1^+^ BMSCs. The differences of targets could potentially be decided by the distinct epigenetic landscape observed in the multiomics data. In the Hmgb2^+^ BMSC, minimal genes and motifs were identified in Pando, indicating a network with *Zeb1*, *Zeb2* and *Klf7* as the regulators (Figure 4E).

In the cultured BMSC, the network is characterized by the prevalence of AP-1 and TEAD factors (Figure 4F). Aside from the highly active AP-1 factors captured based on the motif activity, what was distinct in the regulatory networks was the activated TEAD expression captured in the network. When we narrowed down to the TEAD1 transcription factor and identified all potential downstream targets, we observed that this factor primarily regulated genes related to actin formation, including *Cald1*, *Tagln*, *Actb*, *Actn1*, *Acta2*, *Actg1*, and *Tpm1*, while it suppressed the expression of *Cxcl12* (Figure 4G).

### Hippo Pathway Regulates Multipotency of BMSCs

The upregulation of *Tead1* during early expansion of BMSCs (Figure 4A), along with the subsequent increase of its motif activity in multipotent BMSCs prior to differentiation (Figure 2A), drew our attention. This was because *Tead* TFs and YAP/TAZ co-activators promote proliferation and inhibit differentiation in other stem and progenitor cell populations, including satellite cells of skeletal muscle^47^, neural progenitors^48^, and skin progenitors^49^. Active canonical Hippo signaling limits TEAD activity by phosphorylating transcriptional co-activators YAP/TAZ through LATS kinases and thus preventing their translocation to the nucleus. There was seemingly conflicting data on the role of the Hippo-YAP/TAZ-Tead axis on multipotent BMSC multipotency and differentiation^50^. For example, knockdown of TAZ in murine multipotent BMSCs enhances adipogenic and inhibits osteogenic in vitro differentiation^51^ whereas overexpression of YAP inhibits osteogenic differentiation^52^

Our multiomics data support a model where TEAD inhibits exit from a multipotent state. To test this model, we assessed adipogenic and osteogenic differentiation of multipotent BMSCs pre-treated with small molecule modulators of the Hippo-YAP/TAZ-Tead axis. K975 irreversibly inhibited TEAD from binding to YAP/TAZ^53^. TRULI prevented LATS kinase phosphorylation of YAP/TAZ and inactivated Hippo signaling pathway, resulting in YAP/TAZ translocation to the nucleus (Figure S4A) and constitutive activation^54^. First, we optimized inhibitor concentrations based on colony-forming efficiency (Figure S4B). Next, we measured the extent to which inhibiting and activating TEAD altered the differentiation ability of multipotent BMSCs. During adipogenic differentiation, oil red O staining highlighted an increase in lipid storage, along with an augmentation in lipid vacuole formation within mature adipocytes (Figure 5A). Notably, not only did the number of adipocytes increase, but there was also a concurrent elevation in the expression of the adipocyte marker FABP4 (Figure 5B and Figure 5C). Conversely, activation of YAP/TAZ with TRULI significantly reduced adipogenic differentiation in both number of adipocytes and decrease in the expression of FABP4. In osteogenic differentiation, TEAD inhibition upregulated osteocalcin expression (Figure 5D and Figure 5E) and modestly increased calcium deposition, as shown by alizarin red staining (Figure 5D and Figure 5F). While TRULI pretreatment did not suppress osteocalcin expression, it did markedly reduce calcium deposit, supporting its inhibitory effect on osteogenic differentiation.

**Figure 5:**
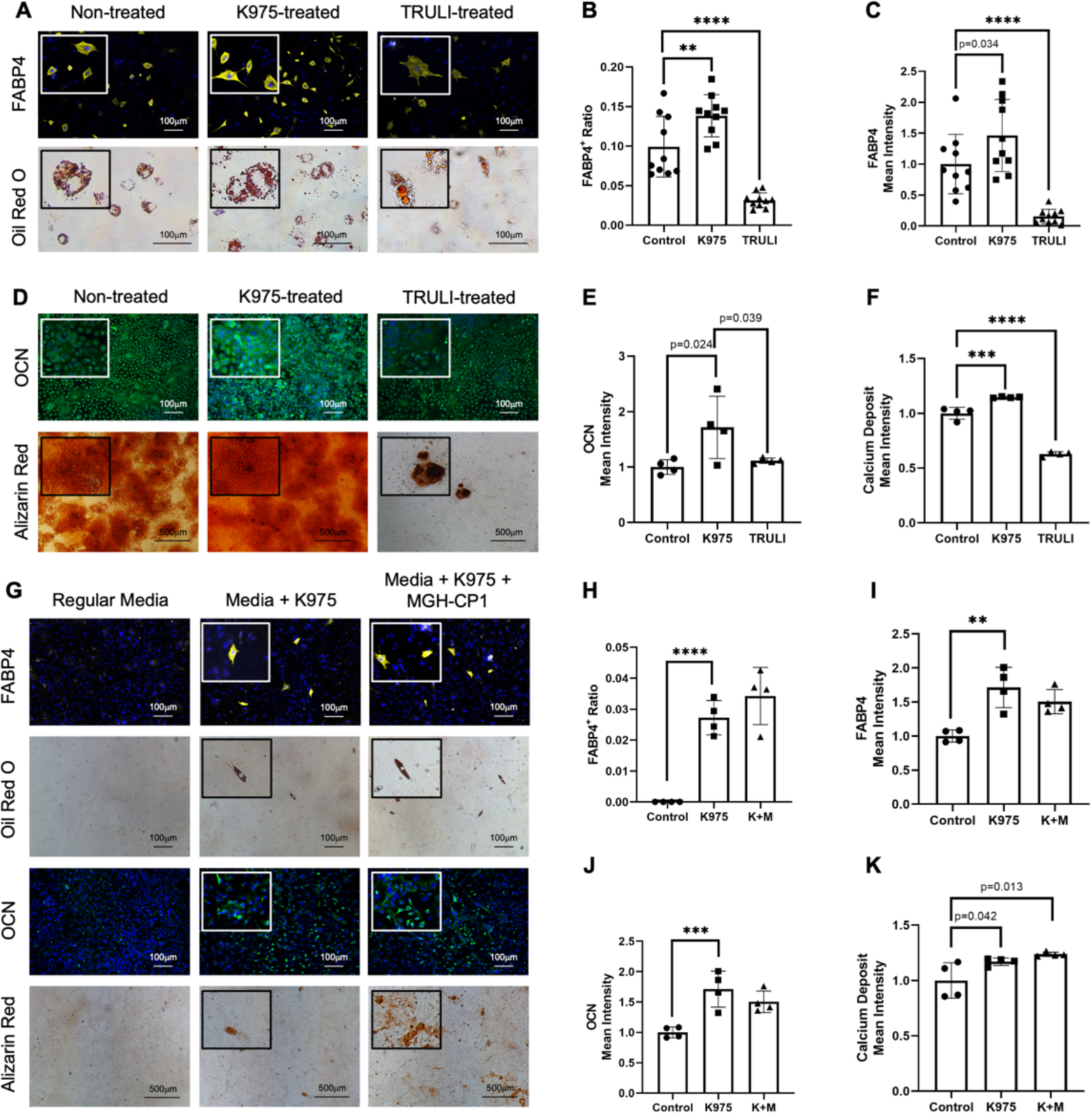
Hippo-Tead Pathway Regulates BMSC Multipotency. (A) Inhibition of the interaction between YAP/TAZ and TEAD with K975 promotes adipogenesis in vitro, while inhibiting YAP/TAZ degradation via the LATS inhibitor TRULI results in the opposite effect. (B) Upregulation in adipogenesis is reflected in increased FABP4 protein expression in K975-pretreated cells (*n* = 10). (C) The percentage of adipocytes over the total number of cells increases upon pretreatment with K975 (*n* = 10). (D) K975 pretreatment promotes osteogenesis *in vitro*, while TRULI pretreatment results in the opposite effect. (E) Osteocalcin expression increases in K975-pretreated BMSCs upon osteogenic differentiation (*n* = 4). (F) Calcium deposits, revealed by Alizarin Red, slightly increase with K975 pretreatment, while TRULI significantly inhibits calcium deposition (*n* = 4). (G) Combined YAP/TAZ-TEAD covalent bond inhibitor K975 and non-covalent bond inhibitor MGH-CP1 drive spontaneous differentiation *in vitro*. (H) FABP4 expression is increased over the baseline background in either K975 or K975+MGH-CP1 groups (*n* = 4). (I) The control group shows no detectable mature adipocytes, while K975 or K975+MGH-CP1 groups reveal a much higher number of mature adipocytes (*n* = 4). (J) Osteocalcin expression increases in either K975 or K975+MGH-CP1 groups (*n* = 4). (K) The control group shows no observable calcium deposit, while K975 and K975+MGH-CP1 groups show detectable Alizarin Red-positive calcium deposits (*n* = 4). (Significant level, **, p < 0.01; ***, p < 0.001, ****, p < 0.0001)

Our experiments targeting the Hippo pathway showed that its inactivation inhibited BMSC differentiation. Next, we aimed to determine the extent to which this inhibitory effect has a lineage bias. To evaluate the impact of Hippo on cell fate decisions, we implemented a regimen to concurrently induce adipogenic and osteogenic differentiation^55^, creating a competitive environment for BMSCs to determine their fate following pretreatment with K975 or TRULI (Figure S4C). While comparing the competitive assay with the uni-lineage differentiation, we observed a similar quantitative trend of promoting adipocyte number (Figure S4E) but a lesser extent of FABP4 expression increase (Figure S4F), comparable osteocalcin expression (Figure S4G), and a significant higher of calcium deposit (Figure S4H). Pretreatment with TRULI revealed a consistent inhibition of BMSCs differentiation across both lineages. These findings indicated that the activation of the Hippo pathway affected differentiation potential without impacting the decision towards either adipogenic or osteogenic fate, consistent with a model where TEAD inhibits an initial exit from a multipotent state that precedes cell fate selection.

To evaluate the efficacy of the Hippo activation in directing multipotent BMSCs to differentiate, we treated these cells with K975 alone or in combination with MGH-CP1, a non-covalent bond TEAD inhibitor ^56^. Our observations revealed that, in the presence of TEAD inhibitors, a subset of BMSCs successfully transitioned from the multipotent state and differentiated into mature adipocytes and osteoblasts. This was evidenced by visible lipid accumulation and calcium deposition (Figure 5G). However, it is noteworthy that the efficiency of this spontaneous differentiation process was notably lower compared to the conventional differentiation regimen. Specifically, the percentage of mature adipocytes constituted less than 5% (Figure 5H).

Additionally, the mean FABP4 expression remained higher than the undifferentiated control (Figure 5I). For osteogenic differentiation, we observed a higher mean osteocalcin intensity under TEAD inhibitors, both with K975 alone and in combination with MGH-CP1 (Figure 5J). This trend was observed similarly in calcium deposits (Figure 5K).

## Discussion

The multipotent BMSCs uncovered from the tri-lineage differentiation experiment exhibited a distinct profile. Although our prediction of multipotency was based on temporal observations, we must acknowledge that a small fraction of the initially expanded cells (14%) is found across other clusters. While it is plausible that the multipotent BMSCs differentiate into downstream intermediate states, we cannot rule out the possibility that this small fraction also expands and contributes to a portion of the differentiated population. To precisely determine the lineage relationship across clusters, barcoding of the original populations is necessary. Moreover, the multipotent BMSCs identified *in vitro* do not correspond to any freshly isolated BMSCs, indicating that the *in vitro* activation alters the cells into a completely different state. This discrepancy can potentially explain the inconsistency in translating some of the *in vitro* results back to *in vivo* conditions.

In contrast to the multipotent BMSCs characterized *in vitro*, computational prediction identified Hmgb2^+^ BMSCs as the most potent stromal cells, a surprising finding given that debates mostly focused on the Cxcl12^+^ BMSCs and osteogenic subclusters. Hmgb2 is known for maintaining the active state in various stem cells, including those of neural^57^, muscle ^58^, hematopoietic^59^, and intestinal^60^ origins. Given its role in regulating cell proliferation, essential for both repopulation and differentiation, it is not surprising that Hmgb2^+^ BMSCs are predicted as the most potent. However, these findings are based purely on computational predictions. To definitively establish whether these populations serve as active stem cells in adult bone marrow stroma, additional studies, potentially involving *in vivo* knock-out or lineage tracing experiments, are warranted. Additionally, the prediction tool “StemFinder” uses cell cycle stochasticity as a reference to assess stemness. This prompts the consideration that Hmgb2^+^ BMSCs may not represent a distinct cell type but rather a transient state of dividing BMSCs from various clusters.

From the lineages inferred by both experimental multi-seq barcoding and computational tools, we observed lineage bifurcations among Limch1^+^ and Col11a1^+^ osteoprogenitor cells as well as between chondroprogenitors and preadipocytes. This finding challenges the theory that endochondral ossification involves chondroprogenitors serving as precursor cells of osteogenic lineage cells. However, since our observations are limited to *in vitro* conditions, we cannot confirm whether the same trajectories are activated during development or under certain pathological conditions. Extending sample collection beyond a week is crucial to fully map the entire trajectory of each lineage. The differences observed in lineage pathways *in vitro* suggest that the behavior of multipotent stem cells *in vitro* differ from their *in vivo* counterparts. Additionally, throughout the differentiation process, we observed a consistent underlying epigenetic state, enriched in AP-1 and TEAD motifs, which persisted despite subculture passages. This persistence underscores the potential of multipotent BMSCs to maintain a degree of lineage flexibility during differentiation until they reach definitive states, such as the adipocyte-like cells, where they become epigenetically locked.

In comparing freshly isolated BMSCs with those differentiating *in vitro*, we observed a consistent differentiation pattern toward states resembling the freshly isolated BMSCs. This observation suggests that the majority of BMSCs *in vivo* may not be in the potent state but are likely in a terminally differentiated state. Despite the success of a chemical regimen in guiding BMSCs towards similar states *in vitro*, we noted discrepancies, such as the absence of mature osteoblastic markers like *Bglap*, *Col1a1*, and *Ibsp* in Limch1^+^ and Col11a1^+^ osteoprogenitor cells, possibly due to the limited incubation time. Moreover, we observed a transient preadipocyte state resembling Cxcl12^+^ BMSCs. Currently, there is no regimen to prevent these cells from further differentiating into adipocyte-like cells *in vitro*. However, TFs inferred from the stabilized Cxcl12^+^ BMSC state, such as *Ebf2* and *Ebf3*, may play essential roles in maintaining these cells. A previous study has shown that *Ebf3* is important to maintain the bone marrow niche^61^. Further comprehensive studies are essential to investigate the factors that contribute to maintaining Cxcl12^+^ BMSCs *in vitro*. Extending the incubation period to capture more mature states of Limch1^+^ and Col11a1^+^ osteoprogenitor cells is crucial for advancing our understanding and techniques in reconstructing the bone marrow niche, both *in vivo* and *in vitro*.

Our network analysis successfully identified key regulators crucial for early transitions in tri-lineage differentiating BMSCs. While *Runx2*^62^, *Sp7*^63^, *Mef2c*^64^ are recognized for their roles in osteogenesis *in vivo*, our *in vitro* finding highlights the importance of additional factors AP-1 and NFKB. Prior research indicates that AP-1 inhibits *Pparg*, promoting osteogenesis^65^, and NFKB prevents dedifferentiation of osteoblasts, an observed phenomenon during bone regeneration in zebrafish^66^. This suggests that the osteogenic differentiation *in vitro* recapitulates to some extent what osteogenesis occurs *in vivo*. As differentiation progressed, we observed an upregulation in *Runx2*^67^, *Runx1*^68^, *Glis3*^69^, and *Mef2a*^70,71^, which reported to play pivotal roles in transitioning osteoprogenitor cells to mature osteoblasts. In adipogenesis, the roles of CEBP and PPAR as adipogenic inducers are well-established^72^. Notably, CEBPB upregulation occurs earlier than that of PPARA and PPARG, aligning with previous studies^73,74^. For the chondrogenic lineage, the well-known chondrogenic factor “*Sox9*” is only detectable at the RNA level but not yet exhibits motif activity, suggesting its role may be more crucial in the later stages of chondrogenic maturation rather than in the early commitment of the multipotent stage. Furthermore, *Ebf1*, a TF enriched in Cxcl12^+^ BMSCs, emerged as a highly active regulators across all lineages, highlighting its critical role in the alternative hematopoietic stem cell (HSC) niche, where it facilitates HSC detachment, leading to reduced quiescence and diminished myeloid output^75^.

Our analysis also suggests AP-1, *Tead1*, and *Zeb2* as potential regulators of multipotency in BMSCs, each noted for their high expression or activity. Although AP-1 factors exhibit high motif activity, we initially did not prioritize investigating AP-1 due to its extensive involvement with many factors, potentially leading to redundancy and posing significant challenges for perturbation^76^. Moreover, complete inhibition of AP-1 factors rapidly induces cell death, hindering further investigation^77^. Despite these challenges, there is evidence suggesting its involvement in regulating stemness. For instance, JUN, an AP-1 factor, exhibits distinct functions in human reprogramming by promoting fibroblast enhancer accessibility and fibroblast-specific gene expression^78^. AP-1 factors also accelerate repopulation of intestinal stem cells post-radiation^79^. In contrast to AP-1, *Zeb2* is active only at the expression level, with low motif activity, as it is one of the downstream targets of Hippo pathway mediators Yap/Taz and *Tead1*^80^. It is likely that *Zeb2* is upregulated by *Tead* regulators, but without significant TF activity.

While the Hippo pathway’s role in regulating BMSC multipotency remains under-explored, it is well known to regulate various stem or progenitor populations in adults. High YAP1 activity promotes proliferation in intestinal stem cell^81,82^, and YAP also promotes the expansion and proliferation of basal epidermal progenitors, inhibiting their terminal differentiation^49,83^. Additionally, YAP’s nuclear translocation promotes the expansion of hepatocytes and oval cells^84^. Existing literature also highlights the impact of Hippo pathway on fate decisions rather than multipotency in BMSCs. For instance, studies have shown that TAZ is crucial in modulating stem cell differentiation^51,85,86^. However, unlike our observation of spontaneous differentiation, TAZ knockout severely impedes the development of the skeletal structure. This suggests that the Hippo pathway is likely involved not only in multipotency maintenance, but also the maturation of the osteogenic lineage. Another study that supports this notion found that YAP/TAZ mediates morphological changes in cells to regulate BMSC commitment to the osteogenic lineage while inhibiting adipogenic and chondrogenic differentiation^87^. Moreover, we have also noticed that *Tead* is upregulated in the osteogenic lineage, even in the absence of an active motif. One study offers a plausible explanation by suggesting that the Vestigial-like family factor VGLL4, predominantly found in the osteoblastic lineage, act as a YAP-TEAD repressor. VGLL4 disrupts the TEAD-mediated transcriptional inhibition of RUNX2, thus promoting osteogenesis by upregulation of RUNX2^88^.

In the *Tead1* regulatory network, we observed that most downstream targets of *Tead1* were associated with cytoskeletal-actin formation, aligning with a model proposed by a study on Ewing Sarcoma ^89^. However, the connection between cytoskeletal genes and the processes of self-renewal and multipotency remains unclear. Despite this, several studies have demonstrated the influence of cytoskeletal dynamics on cell fate decisions. For instance, one study highlighted that TPM1 and YAP play crucial roles in the osteogenesis of BMSCs^90^. However, their influence is primarily observed during differentiation and fate decision processes, rather than in the maintenance of self-renewal or multipotency. It is important to note that the current gene regulatory network reconstruction tools still present significant limitations in accurately predicting appropriate TFs and their targets. Therefore, we cannot rule out the possibility that *Tead1* may also regulate other genes that are more directly linked to BMSC self-renewal and multipotency.

### Concluding Remarks

In this study, we took a single nucleus multiomics approach to identify transcriptional networks that regulate BMSC multipotency *in vitro*. This led to the discovery of the prominent role that the *Tead* family of transcription factors play in regulating the multipotent state of BMSCs. Pre-treatment of BMSCs with TEAD inhibitors promoted adipogenic and osteogenic differentiation, whereas inactivating Hippo, the signaling pathway that controls TEAD activity, via the LATS inhibitor TRULI reduced BMSC differentiation potential. Additionally, we found that TEAD inhibition alone was sufficient to trigger spontaneous differentiation, albeit with lower efficiency than guided differentiation. Collectively, our study sheds light on the complex regulatory mechanisms governing BMSC multipotency and highlights the potential for manipulating BMSC fate in regenerative medicine applications.

## Materials and Method

### Animal and Bone Marrow Tissue Preparation

The animal procedures and care for this study were thoroughly reviewed and approved by the Animal Care and Use Committee (ACUC) at Johns Hopkins University. Fresh bone marrow cells were obtained from 6-10 week-old C57BL/6 mice following ethical guidelines. Euthanasia was performed by CO_2_ exposure, and two femoral bones were harvested and placed in Medium 199 (ThermoFisher Scientific, Cat#12350039) supplemented with 2% fetal bovine serum (FBS, Millipore Sigma, Cat#F0926).

To prepare the bone marrow cells, the soft tissue adhering to the femoral bone was carefully dissected and removed, and the epiphysis was excised. Each femur was flushed with 2.5 ml of Medium 199 + 2% FBS using a 5 ml syringe equipped with a 26G needle to collect bone marrow. The resulting suspension was then pipetted with 1000 μl filtered tips to break down the marrow pieces.

### BMSC Expansion and Subculture

The freshly isolated bone marrow cells were suspended in BMSC culture medium, consisting of ɑMEM, 20% MSC-qualified FBS (Millipore Sigma, Cat#ES-020-B), and 1% PSG (Penicillin, Streptomycin (ThermoFisher Scientific, Cat#15140122), and GlutaMAX (ThermoFisher Scientific, Cat#35050-061)), and seeded in a P100 dish at a density of 0.5 million/cm^2^.

For subculturing the cultured BMSCs, the supernatant culture medium was removed, and the cells were washed once with dPBS (ThermoFisher Scientific, Cat#14190250). Subsequently, the cells were digested with TrypLE for 5 minutes at 37°C (ThermoFisher Scientific, Cat#12605010) and neutralized with supernatant culture medium. The cells were re-suspended in BMSC culture medium and seeded at a density of 3,000-5,000 cells/cm^2^. The BMSC culture medium was changed every three days until the confluency reached 80% for the next subculture.

### *In vitro* BMSC Tri-lineage Differentiation

The BMSC differentiation assays are modified based on previously established protocols^91^.

For adipogenic differentiation, BMSCs were plated at a density of 8,000 cells/cm^2^ in BMSC culture medium and then transitioned to StemPro adipogenic differentiation medium (ThermoFisher Scientific, Cat#A1007001) once they reached confluency.

For osteogenic differentiation, BMSCs were plated at a density of 4,000 cells/cm^2^ in BMSC culture medium. Upon reaching 80% confluency, the BMSC culture medium was replaced with osteogenic differentiation medium. This medium contained 0.1μM Dexamethasone (Millipore Sigma, Cat#D8893), 5mM β-glycerol phosphate (Millipore Sigma, Cat#50020), 100μM Ascorbic acid-2-phosphate (Millipore Sigma, Cat#A8960), and 1% PSG in ɑMEM and 20% MSC-qualified FBS.

For chondrogenic differentiation, 0.5 million BMSCs were centrifuged at 500g for 3 minutes in a 15 ml polypropylene conical tube containing 1 ml of StemXVivo chondrogenic medium (Human/Mouse chondrogenic supplement (R&D, Cat#CCM006) in DMEM/F12 (ThermoFisher Scientific, Cat#11320033). The tube was then loosely capped and left partially unscrewed.

### BMSC Single-cell Preparation

To prepare the fresh bone marrow single-cell suspension, the bone marrow flush was digested with 1 mg/mL STEMxyme1 (Worthington, Cat#LS004106) and 1 mg/mL Dispase II (Millipore Sigma, Cat#D4693) in Medium 199 + 2% FBS for 25 minutes at 37°C in a flask with a stirring bar rotating at 200rpm. Following digestion, the cell suspension was filtered through a 70μm filter (Fisher Scientific, Cat#22-363-548) into a 50ml conical tube.

For single-cell suspension from the dish, cells were digested in 5ml of a 1 mg/mL STEMxyme1 + 1 mg/mL Dispase II solution for 25 minutes at 37°C in the dish. Subsequently, 3ml of TrypLE (ThermoFisher Scientific, Cat#12605010) was added for 5 minutes at 37°C. The reaction was inactivated by adding 10ml fresh BMSC culture medium, followed by a single 10ml dPBS wash. Freshly isolated bone marrow cells and samples from day 1 and day 3 were then stained with conjugated antibodies in Medium 199 supplemented with 2% FBS for 30 minutes on ice for fluorescence-activated cell sorting (FACS).

The antibodies used included TER119-PE (BioLegend, Cat#116208, clone TER-119) for erythrocytes, CD45-PE (eBioscience, Cat#12-0451-82, clone 30-F11) for hematopoietic lineage cells, and CD31-PE (BioLegend, Cat#102507, clone MEC13.3) for endothelial cells. Dead cells and debris were excluded based on FSC, SSC profiles, and PI staining (ThermoFisher Scientific, Cat#P1304MP). FACS was performed using either the MoFlo XDP sorter (Beckman Coulter, Cat#ML99030) or MoFlo Legacy sorter (Beckman Coulter), and the sorted BMSCs were collected in Medium 199 supplemented with 2% FBS.

### Nuclei isolation

The single-cell pellet, containing approximately one million cells, was resuspended in 100μl lysis buffer. The lysis buffer composition included 10mM NaCl (Millipore Sigma, Cat# S8776), 3mM MgCl_2_ (Millipore Sigma, Cat#M1028), 1% BSA (Miltenyi Biotec, Cat#130-091-376), 0.1% Tween-20 (Bio-Rad, Cat#1610781), 1mM DTT (Millipore Sigma, Cat#43816), 1U/μl RNase inhibitor (Millipore Sigma, Cat#3335399001), 0.1% Nonidet P40 Substitute (Millipore Sigma, Cat#98379-10ML-F), and 0.01% Digitonin (ThermoFisher Scientific, Cat#BN2006) in 10mM Tris-HCl buffer (ThermoFisher Scientific, Cat#15567027). This mixture was kept on ice for 5 minutes. Subsequently, 1ml of wash buffer containing 10mM NaCl, 3mM MgCl_2_, 1% BSA, 0.1% Tween-20, 1mM DTT, and 1U/μL RNase inhibitor in 10mM Tris-HCl buffer was added, followed by centrifugation at 500rcf for 5 minutes at 4°C. The wash procedure was repeated for a total of three washes. Finally, the nuclei pellet was resuspended in 20μL of diluted nuclei buffer (10x Genomics, Cat#2000153). Nuclei quality and concentrations were determined using Countess III FL (ThermoFisher Scientific, Cat#AMQAF2000).

### Multi-seq Barcoding

The nuclei suspension underwent transposition following the adjusted 10X protocol. In summary, 18,000 nuclei were suspended in 2.5μL of nuclei buffer and mixed with 5μL of 10X ATAC transposition mix. The mixture was then incubated for 60 minutes at 37°C. Subsequently, the nuclei were suspended in 180μL of cold PBS, and 20μL of 10X anchor-barcode solution was added. The mixture was incubated for 5 minutes on ice before the addition of 20μL co-anchor, followed by an additional 5-minute incubation on ice. Finally, 1ml of cold 1% BSA in PBS was added for two washes. The samples were pooled and re-suspended in 15μL of diluted nuclei.

### Single nucleus multiomics

For each individual sample or multiplexed sample, 18,000 nuclei in 15μL of diluted nuclei were used for barcoding and library preparation. The subsequent procedure was carried out at the Single Cell & Transcriptomics Core at Johns Hopkins University, following the 10X Genomics Chromium Next GEM Single Cell Multiome protocol CG000338 RevE.

### snMultiomics Data Processing

The FASTQ files were processed using Cell Ranger ARC 2.0.2 with default parameters. The pipeline executed alignment, filtering, barcode counting, peak calling, and counting for both ATAC and GEX fractions to generate the BAM file. The resulting BAM files were then utilized to estimate RNA velocity by mapping unspliced and spliced read counts using the Velocyto pipeline.

The aligned RNA data was processed in Scanpy, while the aligned ATAC data was processed in EpiScanpy^92^ to exclude low-quality nuclei. We excluded outliers with more than 20,000 counts or mitochondrial gene expression exceeding 60%. Additionally, cells with ATAC features lower than 500 were removed. Gene counts were normalized to 10,000 per cell, and the most variable genes were identified with a minimal dispersion threshold of 0.5. After cleanup, individual cells were clustered using the Leiden method.

### Differentially Expressed Gene Analysis

Differentially expressed genes (DEGs) were identified for each cell type compared with the remaining clusters using a t-test. The p-values were adjusted using the Benjamini-Hochberg method. Genes with an adjusted p-value < 0.01 were considered statistically significant.

### Cell Cycle Phase Estimation by Tricycle

The cell cycle phase of each cell was estimated using the R package Tricycle v1.0.0^93^. The normalized expression matrix was initially projected into cell cycle space, and subsequently, the cell cycle stages were estimated using the function “estimate_cycle_position”. The “G1/G0” phase was defined between -0.5π and 0.5π, the “S” phase between 0.5π and π, and the “G2/M” phase between π and 1.5π.

### StemFinder

The evaluation of single-cell fate potency was performed using the R package StemFinder (https://github.com/cahanlab/stemfinder). Briefly, a K-nearest neighbor (KNN) graph was constructed based on Euclidean distances in the PCA space. Subsequently, the Gini impurity was computed for each cell, divided by the maximum value, and subtracted from one to obtain the potency score.

### Trajectory inference by RNA velocity

To identify transitions between cell states, we calculated RNA velocity using Velocyto (v0.17)^94^. The raw data were processed through Velocyto to generate a loom file with exon and intron annotations. Subsequently, Velocyto.py was applied to obtain RNA velocity projections onto either UMAP or diffusion maps. In summary, we identified the top 3000 most variable genes with a minimum of 3 counts and detection in 3 cells. Spliced and unspliced counts were normalized separately to the mean total counts. The normalized data underwent Gamma fitting, and RNA velocity was estimated. To determine pseudotime, we defined the origin of RNA velocity as the root and assigned pseudotime using the “tl.dpt” function in Scanpy.

### Gene Enrichment Analysis

The gene enrichment analysis was conducted using the Python package GSEApy^95^. The top 100 differentially expressed genes were imported to assess enrichment in gene sets listed in GO Biological Process 2021, Reactome 2022, and WikiPathway 2021. The most relevant genesets related to stem cell function, cell differentiation, and BMSC-related processes were selected for visualization in the bar chart.

### RNA and ATAC Joint Embedding

The RNA and ATAC layers of snMultiomics data underwent batch correction using the Harmony algorithm^96^. Following this, the graphs produced by the Harmony algorithm were utilized in the Signac package’s FindMultiModalNeighbors function to calculate the joint embedding of individual cells.

### SingleCellNet classification

The classification was performed using singleCellNet^97^ (http://github.com/pcahan1/singleCellNet/) through a two-step protocol involving classifier building and query data classification. In the first step, the classifier was constructed using preprocessed scRNA-seq data with cluster annotation. For each cluster, the top ten most differentially expressed genes were identified, and the top 25 gene pairs were ranked. Based on these selected gene pairs, the preprocessed training data were transformed and utilized to build a random forest classifier consisting of 1000 trees.

In the second step, we classified the query data using the trained classifier to obtain the results. Cells that were not classified as any cell type from the reference data were considered miscellaneous. Clusters that exhibited differential expression of hematopoietic lineage markers (*Ptprc*) and endothelial markers (*Cdh5* and *Pecam1*) were excluded from the analysis.

### Motif Analysis

The motif analysis was conducted using the R tool Signac. The murine Position Frequency Matrix (PFM) was acquired from the precompiled cisBP database^98^. The motif information was integrated into the Seurat object, and chromVAR was employed to infer TF activity^99^. Differentially active motifs were subsequently computed using the “FindMarkers” function.

### Gene Regulatory Network Reconstruction

For the reconstruction of steady-state gene regulatory networks, we utilized Pando^46^. In summary, motif regions overlapping with peaks were connected to nearby genes, and correlations between TFs and their targets, as well as between peaks and their targets, were computed. Subsequently, peaks correlating with target gene expression and TFs associated with motifs in these peaks were selected.

For dynamic gene regulatory network reconstruction, we employed the modified Epoch method^36^. Transcription factors were identified, and gene-gene correlations were calculated. The gene regulatory network was reconstructed based on these correlation values. This step increased the statistical power and eliminated redundant factors crucial to the transition process.

### Colony Forming Efficiency Assay

1.5 million murine cells were plated into a 60 mm diameter Petri dish. After one week of incubation, the cells were fixed with methanol for 30 minutes at room temperature and stained with saturated crystal violet (Millipore Sigma, Cat#C0775). The number of colonies was counted under a dissecting microscope, using a threshold of approximately 50 cells.

### *In vivo* transplantation

The transplantation procedure was conducted following the published protocol^91^ with minor adjustments. Gelfoam collagen sponges (Pfizer, Cat#660-09039604005) were cut into 1cm^2^ square cubes. The sponge pieces were compressed with forceps to remove air and gradually released while seeding 1.5 million cells in 100μL on a Petri dish. The seeded carrier was then incubated for 30 minutes at 37°C.

For preoperative preparation, the mouse was anesthetized with Isoflurane (MWI Veterinary Supply, Cat#502017) using a non-rebreathing circuit (Parkland Scientific, Cat#A518). The dorsal skin surface was shaved and sterilized with betadine and 70% ethanol. Following this, the surgery began with a single 1.5 cm longitudinal incision using sterile scissors. The incision was extended by sterile forceps to accommodate the insertion of seeded Gelfoam transplants. Finally, the incision was closed with two AutoClips (Fine Science Tools, Cat#12020) and tissue glue (3M, Cat#B004C12Q46).

### Faxitron X-ray Imaging

The X-ray image was captured using the Faxitron MX-20 system (Faxitron, Cat#XRAY82033-2263) to visualize the subcutaneous transplant. The default power and exposure were initially set to 18kV for 10 seconds. The voltage was then adjusted to obtain an image with optimized exposure.

### Histological Preparation

The transplant was harvested by tissue dissection, and the *in vitro* chondrogenic differentiation pellet was washed with dPBS. The sample was then fixed overnight in 10% Formalin (Millipore Sigma, Cat#HT501128). For frozen sections, the fixed samples were processed at the Reference Histology core at Johns Hopkins University. In summary, the samples were dehydrated in 30% sucrose in PBS overnight, embedded in O.C.T compound (Ted Pella, Cat#27050), and sectioned at 10μm. For paraffin sections, the fixed tissue was embedded in paraffin and sectioned at 5μm.

### Oil Red O Staining

Cells in the dish are washed with PBS and subsequently fixed in 10% formalin for 30 minutes at room temperature. After fixation, cells in the dish or the slides from the frozen section were rinsed with deionized (DI) water followed by 60% isopropanol. A stock solution of 0.5% oil red O (Millipore Sigma, Cat#O1391) in isopropanol (Millipore Sigma, Cat#I9516) was then diluted at a ratio of 60:40 in water and added to the fixed cells for 5 minutes at room temperature. Subsequently, the dish or the slide was rinsed with DI water.

### Alizarin Red Staining

The fixed cells in the dish are rinsed with distilled water and stained with Alizarin Red Solution for 30 seconds to 5 minutes. The reaction can be observed with the naked eye, and typically, 2 minutes will produce a nice red-orange staining of calcium. After staining, excess dye is shaken off, and the cells are washed with distilled water three times.

### Alcian Blue Staining

The fixed cells are rinsed three times with PBS. Subsequently, the cells are stained with the Alcian Blue solution (Alcian Blue 1g in 3% Acetic acid 100mL) for 30 minutes. Following staining, the cells are rinsed with distilled water three times.

### Immunofluorescence Staining

The cells in the dish were fixed in 10% buffered formalin for 30 minutes at room temperature, washed with distilled water, and incubated with primary antibodies in antibody diluent at 4°C overnight, followed by three washes in TBST. Subsequently, the dish was incubated with secondary antibodies conjugated with fluorescence at room temperature for 1 hour while avoiding light, followed by three washes in TBST. Nuclear staining was performed with 1:1000 dilution of DAPI in PBS. Images were captured using a Nikon Eclipse Ti-S microscope with DS-U3 and DS-Qi2.

### Fluorescence In Situ Hybridization

Dehydration of slides was performed in a series of ethanol solutions: slides were placed in 70% EtOH for 2 minutes, followed by transfer to 85% EtOH for an additional 2 minutes, and finally transferred to 100% EtOH for another 2 minutes. Subsequently, the slides were allowed to air dry. During the drying period, a probe mixture was prepared for application to the slides. The recommended mixture consisted of 2μL of probe and 8μL of buffer, totaling 10μL per cellular area. Using a pipette, 10μL of the probe mixture was dispensed onto each cellular area, and a 22×22 mm cover glass was carefully placed over the area, ensuring the absence of air bubbles.

The cover glass was sealed with rubber cement, ensuring complete sealing along all edges to prevent drying of the probe under the coverslip. To denature the slides, a hot plate was used; for FFPE tissue slide preparations, the slide was placed on the hot plate at 75°C, covered to shield it from direct light, and kept on the hot plate for 7 minutes. Following this, the slide was removed, placed in a humidified chamber, and then incubated at 37°C for at least 16 hours.

### Hippo modulation experiments

For the pretreatment experiments, expanded cells were treated with a 10μM concentration of the selected TEAD inhibitors (K975 or MGH-CP1) to simulate active Hippo signaling or TRULI to mimic its inactivation, for a duration of three days. Subsequently, the cells were washed with dPBS and incubated with differentiation media. In the spontaneous differentiation experiment, expanded cells were incubated with either 5μM K975 or a combination of 5μM K975 and 5μM MGH-CP1 for at least two weeks. The cultured cells were then washed with PBS and fixed in formalin for subsequent histological analysis.

### Statistical tests

All quantitative measurements were expressed as mean ± standard deviation (s.d.). Statistical comparisons between two distinct groups were conducted using an unpaired Student’s t-test. The level of significance was set at p < 0.05.

## Code and Data Availability

All the FASTQ files have been deposited in the GEO database under the accession number GSE234540 (reviewer password: cbmtummcfxwzfax). The processed data have been uploaded to Amazon AWS and are accessible to the public. The codes required for reproducing the entire analysis can be found on GitHub at: https://github.com/CahanLab/BMSC_23

## Author Contributions

Y.C and P.C conceived the project and wrote the manuscript. C.C and Q.B provided experimental assistance. R.M, S.K and P.R provided various methods and critiqued the manuscript.

## Acknowledgments

P.C received support through grants from R35GM124725 NIH/NIGMS and the Johns Hopkins Catalyst Award. R.M, S.K, and P.R received support through grants from NIDCR ZIA DE000380. We express our gratitude to Dr. Renyuan Bai for supplying Gelfoam and to Dr. Christopher S. McGinnis for providing the Multi-seq reagent prior to its market launch. Special thanks go to Linda D. Orzolek and Tyler J. Creamer from the Johns Hopkins Single-cell & Transcriptomics Core for their expertise in conducting single-cell RNA-sequencing. Additionally, we acknowledge Hao Zhang from the Johns Hopkins Bloomberg Flow Cytometry and Immunology Core for his contribution to cell sorting.

## Supplementary

**Figure S1:**
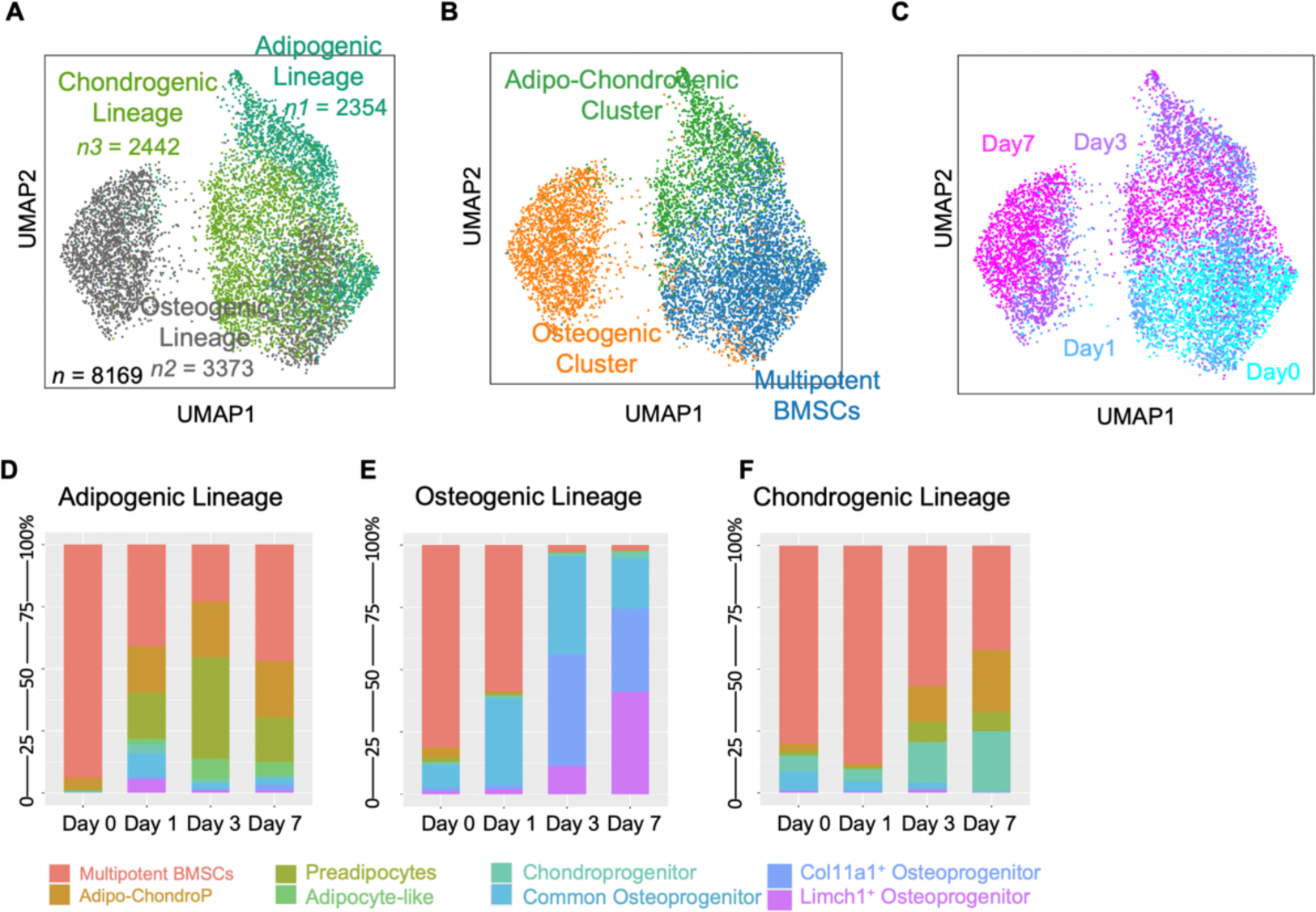
Lineage Specification of Tri-lineage Differentiation of Multipotent BMSCs. (A) Each lineage contributed a similar portion of approximately 3000 cells. (B) Three major clusters detected in the differentiating multipotent BMSCs. (C) Time labels delineate the sequential progression of clusters from day 0 through day 7. (D) Attribution of clusters within the adipogenic lineage to different time points. (E) Attribution of clusters within the osteogenic lineage to different time points. (F) Attribution of clusters within the chondrogenic lineage to different time points.

**Figure S2:**
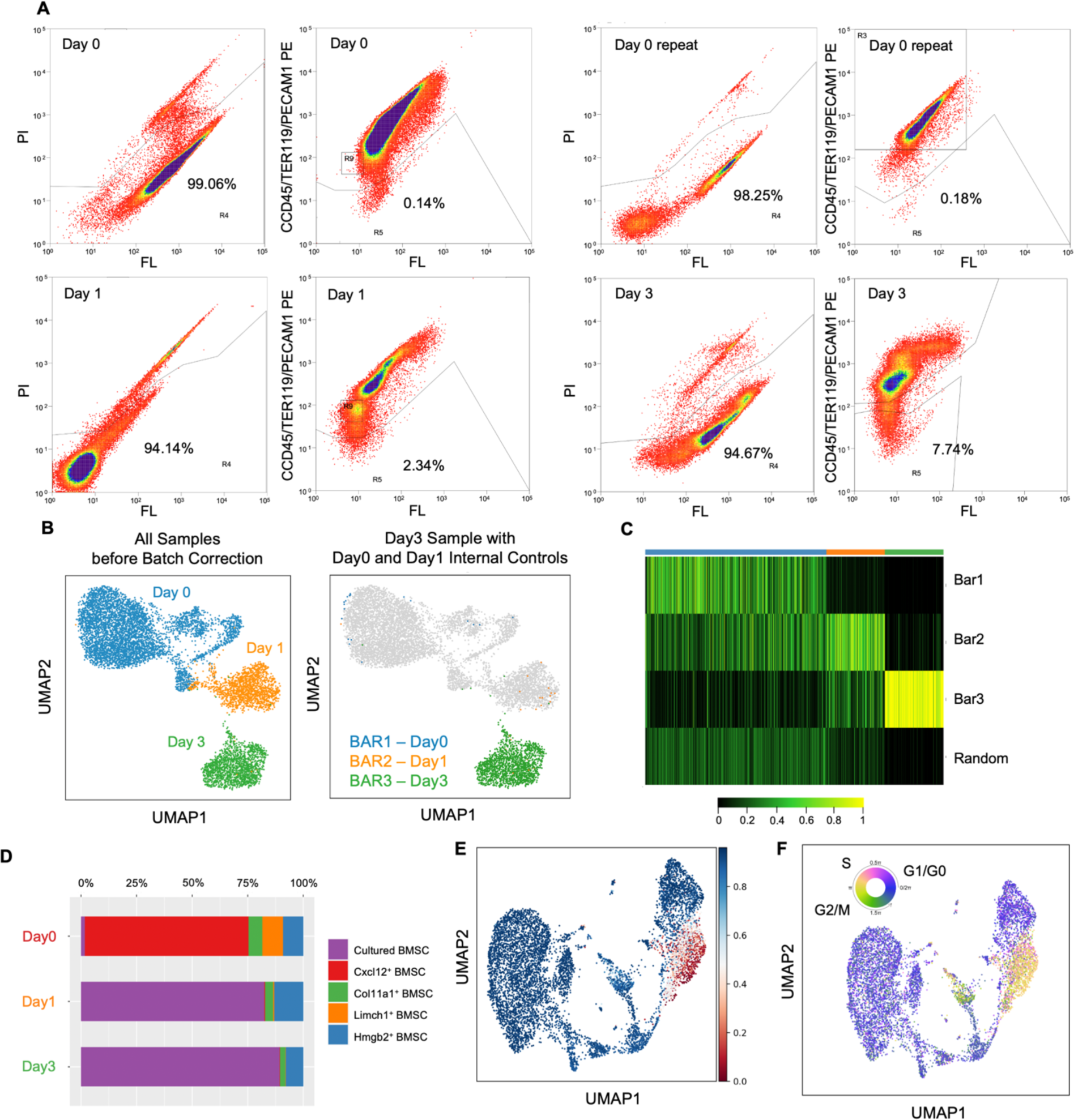
Rapid Loss of Heterogeneity in BMSC Post-isolation. (A) BMSC samples were enriched using FACS to remove confounding cells like hematopoietic and endothelial cells. (B) BMSCs collected from three distinct timepoints exhibit unique features that are visually apparent on UMAP without undergoing batch correction. (C) For the day 3 sample, we utilized Multi-seq techniques by multiplexing Day 0 and Day 1 to act as an internal control. This enabled accurate classification of cells from Day 0, Day 1, and Day 3 to their respective timepoints. (D) The attribution plot illustrates how different BMSC clusters are distributed across various timepoints. (E) StemFinder predicts the basal states of individual cells. (F) Cell cycle analysis suggests that both Hmgb2^+^ BMSCs and cultured BMSCs are in highly active proliferative states.

**Figure S3:**
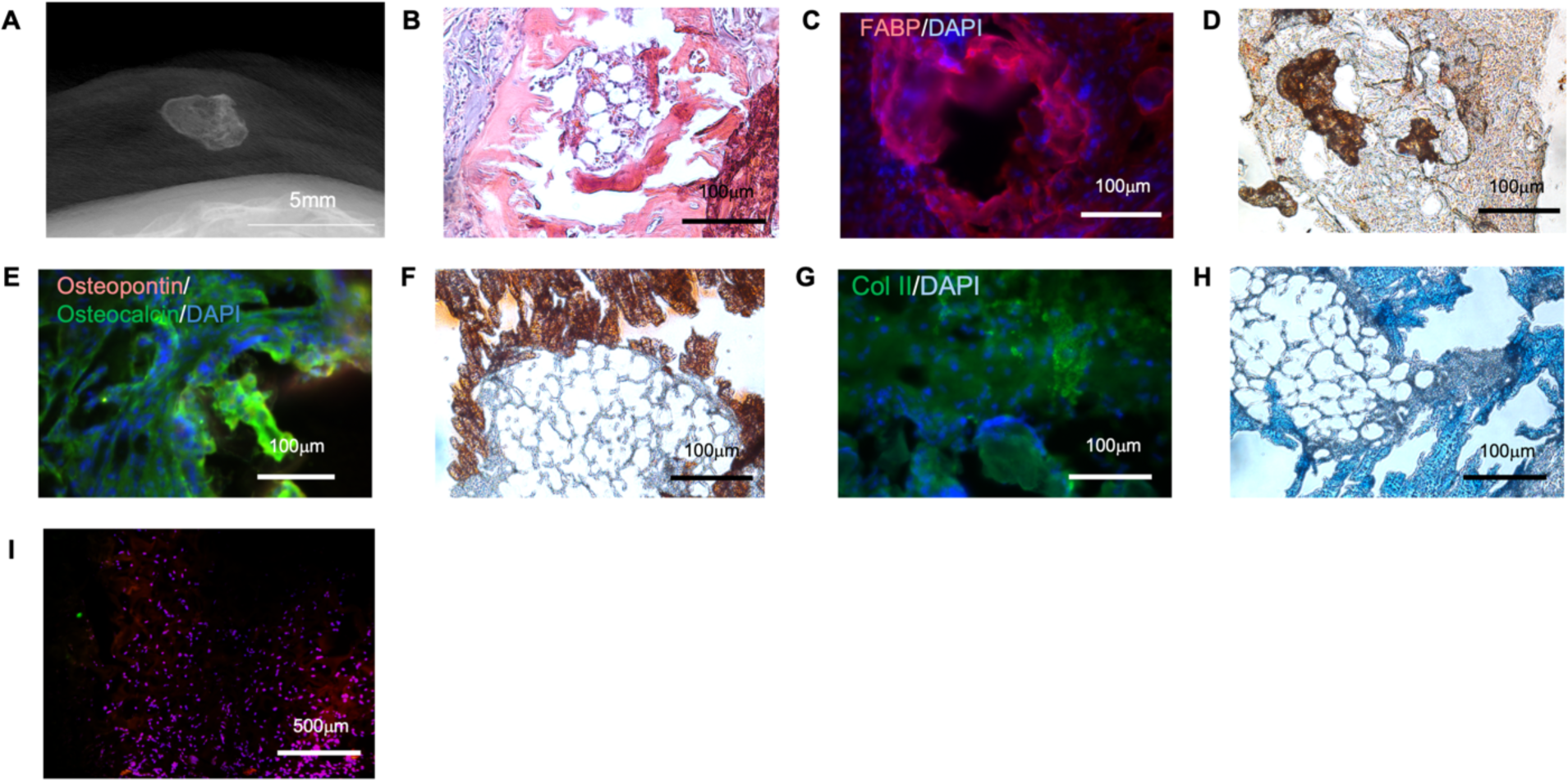
Cultured BMSCs Reconstituted Bone Marrow *in vivo*. (A) Transplantation of cultured BMSCs *in vivo* leads to the spontaneous formation of ossicles and bone marrow. (B) HE staining of the harvested BMSC transplant. (C) Immunohistochemistry staining for FABP in the transplant. (D) Oil Red O staining of the transplant for lipid detection. (E) Immunohistochemical staining for osteocalcin of the transplant, indicating bone formation. (F) Alizarin Red staining reveals positive calcium deposition, indicative of mineralization. (G) Immunohistochemistry staining for type II collagen, suggesting cartilage formation (H) Alcian Blue staining for cartilage detection (I) FISH staining revealed that most of the cells present in the transplant carrier originated from the male donor mouse, as confirmed by Y chromosome painting.

**Figure S4:**
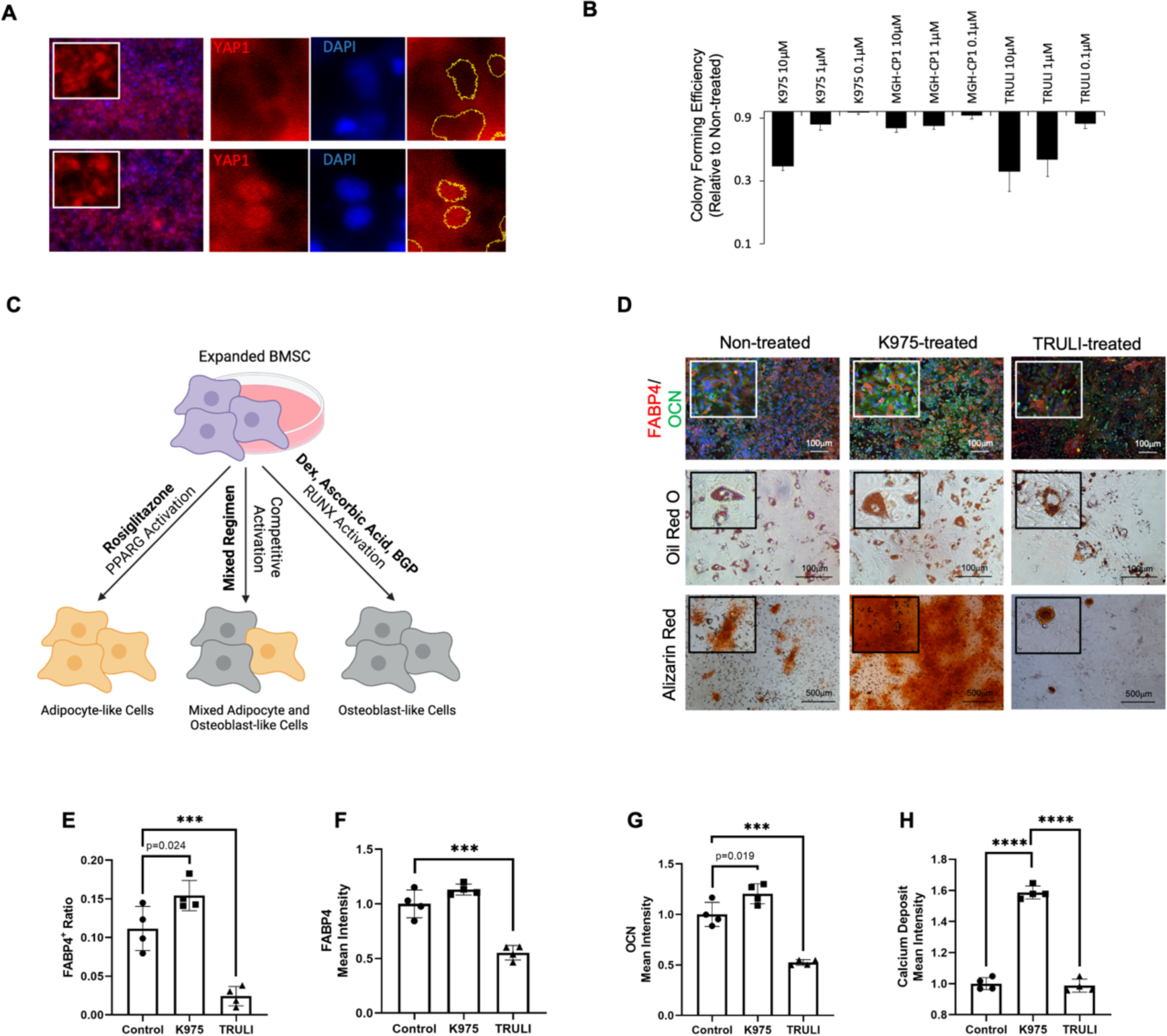
Effect of Hippo-pathway on BMSC Early Fate Decision. (A) Treatment with the LATS inhibitor TRULI results in the translocation of YAP from the cytoplasm to the nucleus under confluent conditions. (B) A colony-forming efficiency assay was conducted to assess cell survival upon inhibitor treatment. (C) Simultaneous adipogenic and osteogenic differentiation of cultured BMSCs was induced using a mixed regimen comprising two independent differentiation protocols. (D) A competitive lineage differentiation assay reveals no lineage preference upon pretreatment with either K975 or TRULI. (E) FABP4 expression decreases with TRULI pretreatment, while K975 pretreatment shows a slight increase without reaching statistical significance (*n* = 4). (F) The percentage of mature adipocytes slightly increases with K975 pretreatment (*n* = 4). (G) Osteocalcin expression increases with K975 pretreatment and decreases with TRULI pretreatment (*n* = 4). (H) Calcium deposition increases with K975 pretreatment, while TRULI pretreatment shows a comparable deposit level as the control without any inhibitor pretreatment (*n* = 4).

## References

1. Bianco, P. & Robey, P. G. Skeletal stem cells. Development 142, 1023–1027 (2015).

2. Jeffery, E. C., Mann, T. L. A., Pool, J. A., Zhao, Z. & Morrison, S. J. Bone marrow and periosteal skeletal stem/progenitor cells make distinct contributions to bone maintenance and repair. Cell Stem Cell 29, 1547–1561.e6 (2022).

3. Veis, D. J. & O’Brien, C. A. Osteoclasts, master sculptors of bone. Annu. Rev. Pathol. 18, 257–281 (2023).

4. Kfoury, Y. & Scadden, D. T. Mesenchymal cell contributions to the stem cell niche. Cell Stem Cell 16, 239–253 (2015).

5. Friedenstein, A. J., Latzinik, N. W., Grosheva, A. G. & Gorskaya, U. F. Marrow microenvironment transfer by heterotopic transplantation of freshly isolated and cultured cells in porous sponges. Exp. Hematol. 10, 217–227 (1982).

6. Shen, B. et al. A mechanosensitive peri-arteriolar niche for osteogenesis and lymphopoiesis. Nature 591, 438–444 (2021).

7. Greenbaum, A. et al. CXCL12 in early mesenchymal progenitors is required for haematopoietic stem-cell maintenance. Nature 495, 227–230 (2013).

8. Sugiyama, T., Kohara, H., Noda, M. & Nagasawa, T. Maintenance of the hematopoietic stem cell pool by CXCL12-CXCR4 chemokine signaling in bone marrow stromal cell niches. Immunity 25, 977–988 (2006).

9. Zhou, B. O., Yue, R., Murphy, M. M., Peyer, J. G. & Morrison, S. J. Leptin-receptor-expressing mesenchymal stromal cells represent the main source of bone formed by adult bone marrow. Cell Stem Cell 15, 154–168 (2014).

10. Kunisaki, Y. et al. Arteriolar niches maintain haematopoietic stem cell quiescence. Nature 502, 637–643 (2013).

11. Pinho, S. et al. PDGFRα and CD51 mark human nestin+ sphere-forming mesenchymal stem cells capable of hematopoietic progenitor cell expansion. J. Exp. Med. 210, 1351– 1367 (2013).

12. Tormin, A. et al. CD146 expression on primary nonhematopoietic bone marrow stem cells is correlated with in situ localization. Blood 117, 5067–5077 (2011).

13. Robey, P. G., Kuznetsov, S. A., Riminucci, M. & Bianco, P. Bone marrow stromal cell assays: in vitro and in vivo. Methods Mol. Biol. 1130, 279–293 (2014).

14. Friedenstein, A. J., Petrakova, K. V., Kurolesova, A. I. & Frolova, G. P. Heterotopic of bone marrow. Analysis of precursor cells for osteogenic and hematopoietic tissues. Transplantation 6, 230–247 (1968).

15. Friedenstein, A. J., Gorskaja, J. F. & Kulagina, N. N. Fibroblast precursors in normal and irradiated mouse hematopoietic organs. Exp. Hematol. 4, 267–274 (1976).

16. Compston, J. E., McClung, M. R. & Leslie, W. D. Osteoporosis. Lancet 393, 364–376 (2019).

17. Ebeling, P. R. Clinical practice. Osteoporosis in men. N. Engl. J. Med. 358, 1474–1482 (2008).

18. Black, D. M. & Rosen, C. J. Postmenopausal Osteoporosis. N. Engl. J. Med. 374, 2096– 2097 (2016).

19. Chawla, A., Schwarz, E. J., Dimaculangan, D. D. & Lazar, M. A. Peroxisome proliferator-activated receptor (PPAR) gamma: adipose-predominant expression and induction early in adipocyte differentiation. Endocrinology 135, 798–800 (1994).

20. Benvenuti, S. et al. Rosiglitazone stimulates adipogenesis and decreases osteoblastogenesis in human mesenchymal stem cells. J. Endocrinol. Invest. 30, RC26-30 (2007).

21. Rosen, E. D. & Spiegelman, B. M. Molecular regulation of adipogenesis. Annu. Rev. Cell Dev. Biol. 16, 145–171 (2000).

22. Gregoire, F. M., Smas, C. M. & Sul, H. S. Understanding adipocyte differentiation. Physiol. Rev. 78, 783–809 (1998).

23. Goldring, M. B., Tsuchimochi, K. & Ijiri, K. The control of chondrogenesis. J. Cell. Biochem. 97, 33–44 (2006).

24. Johnstone, B., Hering, T. M., Caplan, A. I., Goldberg, V. M. & Yoo, J. U. In vitro chondrogenesis of bone marrow-derived mesenchymal progenitor cells. Exp. Cell Res. 238, 265–272 (1998).

25. Reddi, A. H. & Cunningham, N. S. Initiation and promotion of bone differentiation by bone morphogenetic proteins. J. Bone Miner. Res. 8 **Suppl 2**, S499–502 (1993).

26. Kurihara, N. et al. Effects of 1,25-dihydroxyvitamin D3 on osteoblastic MC3T3-E1 cells. Endocrinology 118, 940–947 (1986).

27. Harada, S. & Rodan, G. A. Control of osteoblast function and regulation of bone mass. Nature 423, 349–355 (2003).

28. Franceschi, R. T., Iyer, B. S. & Cui, Y. Effects of ascorbic acid on collagen matrix formation and osteoblast differentiation in murine MC3T3-E1 cells. J. Bone Miner. Res. 9, 843–854 (1994).

29. Marie, P. J. Transcription factors controlling osteoblastogenesis. Arch. Biochem. Biophys. 473, 98–105 (2008).

30. Gadi, J. et al. The transcription factor protein Sox11 enhances early osteoblast differentiation by facilitating proliferation and the survival of mesenchymal and osteoblast progenitors. J. Biol. Chem. 288, 25400–25413 (2013).

31. Larson, B. L., Ylostalo, J., Lee, R. H., Gregory, C. & Prockop, D. J. Sox11 is expressed in early progenitor human multipotent stromal cells and decreases with extensive expansion of the cells. Tissue Eng. Part A 16, 3385–3394 (2010).

32. Sipp, D., Robey, P. G. & Turner, L. Clear up this stem-cell mess. Nature 561, 455–457 (2018).

33. Soliman, H. et al. Multipotent stromal cells: One name, multiple identities. Cell Stem Cell 28, 1690–1707 (2021).

34. Hurley, K. et al. Reconstructed Single-Cell Fate Trajectories Define Lineage Plasticity Windows during Differentiation of Human PSC-Derived Distal Lung Progenitors. Cell Stem Cell 26, 593–608.e8 (2020).

35. Fletcher, R. B. et al. Deconstructing Olfactory Stem Cell Trajectories at Single-Cell Resolution. Cell Stem Cell 20, 817–830.e8 (2017).

36. Su, E. Y., Spangler, A., Bian, Q., Kasamoto, J. Y. & Cahan, P. Reconstruction of dynamic regulatory networks reveals signaling-induced topology changes associated with germ layer specification. Stem Cell Reports 17, 427–442 (2022).

37. Itoh-Nakadai, A. et al. A Bach2-Cebp Gene Regulatory Network for the Commitment of Multipotent Hematopoietic Progenitors. Cell Rep. 18, 2401–2414 (2017).

38. Zhao, Q. et al. TCF21 and AP-1 interact through epigenetic modifications to regulate coronary artery disease gene expression. Genome Med. 11, 23 (2019).

39. Wu, H. et al. Genomic occupancy of Runx2 with global expression profiling identifies a novel dimension to control of osteoblastogenesis. Genome Biol. 15, R52 (2014).

40. Chan, S. F. et al. Transcriptional profiling of MEF2-regulated genes in human neural progenitor cells derived from embryonic stem cells. Genom. Data 3, 24–27 (2015).

41. Obier, N. et al. Cooperative binding of AP-1 and TEAD4 modulates the balance between vascular smooth muscle and hemogenic cell fate. Development 143, 4324–4340 (2016).

42. Lefterova, M. I. et al. PPARgamma and C/EBP factors orchestrate adipocyte biology via adjacent binding on a genome-wide scale. Genes Dev. 22, 2941–2952 (2008).

43. Nielsen, R. et al. Genome-wide profiling of PPARgamma:RXR and RNA polymerase II occupancy reveals temporal activation of distinct metabolic pathways and changes in RXR dimer composition during adipogenesis. Genes Dev. 22, 2953–2967 (2008).

44. Yan, J. et al. Smad4 deficiency impairs chondrocyte hypertrophy via the Runx2 transcription factor in mouse skeletal development. J. Biol. Chem. 293, 9162–9175 (2018).

45. Hoshiba, T., Kawazoe, N., Tateishi, T. & Chen, G. Development of stepwise osteogenesis-mimicking matrices for the regulation of mesenchymal stem cell functions. J. Biol. Chem. 284, 31164–31173 (2009).

46. Fleck, J. S. et al. Inferring and perturbing cell fate regulomes in human brain organoids. Nature 621, 365–372 (2023).

47. Judson, R. N. et al. The Hippo pathway member Yap plays a key role in influencing fate decisions in muscle satellite cells. J. Cell Sci. 125, 6009–6019 (2012).

48. Cao, X., Pfaff, S. L. & Gage, F. H. YAP regulates neural progenitor cell number via the TEA domain transcription factor. Genes Dev. 22, 3320–3334 (2008).

49. Zhang, H., Pasolli, H. A. & Fuchs, E. Yes-associated protein (YAP) transcriptional coactivator functions in balancing growth and differentiation in skin. Proc Natl Acad Sci USA 108, 2270–2275 (2011).

50. Kegelman, C. D., Collins, J. M., Nijsure, M. P., Eastburn, E. A. & Boerckel, J. D. Gone caving: roles of the transcriptional regulators YAP and TAZ in skeletal development. Curr. Osteoporos. Rep. 18, 526–540 (2020).

51. Hong, J.-H. et al. TAZ, a transcriptional modulator of mesenchymal stem cell differentiation. Science 309, 1074–1078 (2005).

52. Sen, B. et al. Intranuclear actin regulates osteogenesis. Stem Cells 33, 3065–3076 (2015).

53. Kaneda, A. et al. The novel potent TEAD inhibitor, K-975, inhibits YAP1/TAZ-TEAD protein-protein interactions and exerts an anti-tumor effect on malignant pleural mesothelioma. Am. J. Cancer Res. 10, 4399–4415 (2020).

54. Kastan, N. et al. Small-molecule inhibition of Lats kinases may promote Yap-dependent proliferation in postmitotic mammalian tissues. Nat. Commun. 12, 3100 (2021).

55. McBeath, R., Pirone, D. M., Nelson, C. M., Bhadriraju, K. & Chen, C. S. Cell shape, cytoskeletal tension, and RhoA regulate stem cell lineage commitment. Dev. Cell 6, 483– 495 (2004).

56. Sun, Y. et al. Pharmacological blockade of TEAD-YAP reveals its therapeutic limitation in cancer cells. Nat. Commun. 13, 6744 (2022).

57. Kimura, A., Matsuda, T., Sakai, A., Murao, N. & Nakashima, K. HMGB2 expression is associated with transition from a quiescent to an activated state of adult neural stem cells. Dev. Dyn. 247, 229–238 (2018).

58. Zhou, X. et al. HMGB2 regulates satellite-cell-mediated skeletal muscle regeneration through IGF2BP2. J. Cell Sci. 129, 4305–4316 (2016).

59. Zhang, C. et al. Latexin regulation by HMGB2 is required for hematopoietic stem cell maintenance. Haematologica 105, 573–584 (2020).

60. Yun, J. et al. Senescent cells perturb intestinal stem cell differentiation through Ptk7 induced noncanonical Wnt and YAP signaling. Nat. Commun. 14, 156 (2023).

61. Seike, M., Omatsu, Y., Watanabe, H., Kondoh, G. & Nagasawa, T. Stem cell niche-specific Ebf3 maintains the bone marrow cavity. Genes Dev. 32, 359–372 (2018).

62. Schroeder, T. M., Jensen, E. D. & Westendorf, J. J. Runx2: a master organizer of gene transcription in developing and maturing osteoblasts. Birth Defects Res. C Embryo Today 75, 213–225 (2005).

63. Zhang, C. Transcriptional regulation of bone formation by the osteoblast-specific transcription factor Osx. J. Orthop. Surg. Res. 5, 37 (2010).

64. Kawane, T. et al. Dlx5 and mef2 regulate a novel runx2 enhancer for osteoblast-specific expression. J. Bone Miner. Res. 29, 1960–1969 (2014).

65. Luther, J. et al. Fra-2/AP-1 controls adipocyte differentiation and survival by regulating PPARγ and hypoxia. Cell Death Differ. 21, 655–664 (2014).

66. Mishra, R., Sehring, I., Cederlund, M., Mulaw, M. & Weidinger, G. NF-κB Signaling Negatively Regulates Osteoblast Dedifferentiation during Zebrafish Bone Regeneration. Dev. Cell 52, 167–182.e7 (2020).

67. Franceschi, R. T. & Xiao, G. Regulation of the osteoblast-specific transcription factor, Runx2: responsiveness to multiple signal transduction pathways. J. Cell. Biochem. 88, 446–454 (2003).

68. Tang, C.-Y. et al. Runx1 is a central regulator of osteogenesis for bone homeostasis by orchestrating BMP and WNT signaling pathways. PLoS Genet. 17, e1009233 (2021).

69. Beak, J. Y., Kang, H. S., Kim, Y.-S. & Jetten, A. M. Krüppel-like zinc finger protein Glis3 promotes osteoblast differentiation by regulating FGF18 expression. J. Bone Miner. Res. 22, 1234–1244 (2007).

70. Wang, C. et al. Lineage-selective super enhancers mediate core regulatory circuitry during adipogenic and osteogenic differentiation of human mesenchymal stem cells. Cell Death Dis. 13, 866 (2022).

71. Rauch, A. et al. Osteogenesis depends on commissioning of a network of stem cell transcription factors that act as repressors of adipogenesis. Nat. Genet. 51, 716–727 (2019).

72. Madsen, M. S., Siersbæk, R., Boergesen, M., Nielsen, R. & Mandrup, S. Peroxisome proliferator-activated receptor γ and C/EBPα synergistically activate key metabolic adipocyte genes by assisted loading. Mol. Cell. Biol. 34, 939–954 (2014).

73. Guo, L., Li, X. & Tang, Q.-Q. Transcriptional regulation of adipocyte differentiation: a central role for CCAAT/enhancer-binding protein (C/EBP) β. J. Biol. Chem. 290, 755–761 (2015).

74. Rosen, E. D. et al. C/EBPalpha induces adipogenesis through PPARgamma: a unified pathway. Genes Dev. 16, 22–26 (2002).

75. Derecka, M. et al. EBF1-deficient bone marrow stroma elicits persistent changes in HSC potential. Nat. Immunol. 21, 261–273 (2020).

76. Wang, W.-M., Wu, S.-Y., Lee, A.-Y. & Chiang, C.-M. Binding site specificity and factor redundancy in activator protein-1-driven human papillomavirus chromatin-dependent transcription. J. Biol. Chem. 286, 40974–40986 (2011).

77. Ye, N., Ding, Y., Wild, C., Shen, Q. & Zhou, J. Small molecule inhibitors targeting activator protein 1 (AP-1). J. Med. Chem. 57, 6930–6948 (2014).

78. Markov, G. J. et al. AP-1 is a temporally regulated dual gatekeeper of reprogramming to pluripotency. Proc Natl Acad Sci USA 118, (2021).

79. Chen, F. et al. TIGAR/AP-1 axis accelerates the division of Lgr5-reserve intestinal stem cells to reestablish intestinal architecture after lethal radiation. Cell Death Dis. 11, 501 (2020).

80. Deng, Y. et al. A reciprocal regulatory loop between TAZ/YAP and G-protein Gαs regulates Schwann cell proliferation and myelination. Nat. Commun. 8, 15161 (2017).

81. Camargo, F. D. et al. YAP1 increases organ size and expands undifferentiated progenitor cells. Curr. Biol. 17, 2054–2060 (2007).

82. Zhou, D. et al. Mst1 and Mst2 protein kinases restrain intestinal stem cell proliferation and colonic tumorigenesis by inhibition of Yes-associated protein (Yap) overabundance. Proc Natl Acad Sci USA 108, E1312–20 (2011).

83. Schlegelmilch, K. et al. Yap1 acts downstream of α-catenin to control epidermal proliferation. Cell 144, 782–795 (2011).

84. Lu, L. et al. Hippo signaling is a potent in vivo growth and tumor suppressor pathway in the mammalian liver. Proc Natl Acad Sci USA 107, 1437–1442 (2010).

85. Tang, Y. & Weiss, S. J. Snail/Slug-YAP/TAZ complexes cooperatively regulate mesenchymal stem cell function and bone formation. Cell Cycle 16, 399–405 (2017).

86. Zhao, X. et al. Yap and Taz promote osteogenesis and prevent chondrogenesis in neural crest cells in vitro and in vivo. Sci. Signal. 15, eabn9009 (2022).

87. Tang, Y. et al. MT1-MMP-dependent control of skeletal stem cell commitment via a β1-integrin/YAP/TAZ signaling axis. Dev. Cell 25, 402–416 (2013).

88. Suo, J. et al. VGLL4 promotes osteoblast differentiation by antagonizing TEADs-inhibited Runx2 transcription. Sci. Adv. 6, (2020).

89. Katschnig, A. M. et al. EWS-FLI1 perturbs MRTFB/YAP-1/TEAD target gene regulation inhibiting cytoskeletal autoregulatory feedback in Ewing sarcoma. Oncogene 36, 5995– 6005 (2017).

90. Brielle, S. et al. Delineating the heterogeneity of matrix-directed differentiation toward soft and stiff tissue lineages via single-cell profiling. Proc Natl Acad Sci USA 118, (2021).

91. Robey, P. G., Kuznetsov, S. A., Bianco, P. & Riminucci, M. Bone marrow stromal cell assays: in vitro and in vivo. Methods Mol. Biol. 2230, 379–396 (2021).

92. Danese, A. et al. EpiScanpy: integrated single-cell epigenomic analysis. Nat. Commun. 12, 5228 (2021).

93. Zheng, S. C. et al. Universal prediction of cell-cycle position using transfer learning. Genome Biol. 23, 41 (2022).

94. La Manno, G. et al. RNA velocity of single cells. Nature 560, 494–498 (2018).

95. Fang, Z., Liu, X. & Peltz, G. GSEApy: a comprehensive package for performing gene set enrichment analysis in Python. Bioinformatics 39, (2023).

96. Korsunsky, I. et al. Fast, sensitive and accurate integration of single-cell data with Harmony. Nat. Methods 16, 1289–1296 (2019).

97. Tan, Y. & Cahan, P. SingleCellNet: a computational tool to classify single cell RNA-Seq data across platforms and across species. BioRxiv (2018) doi:10.1101/508085.

98. Kartha, V. K. et al. Functional inference of gene regulation using single-cell multi-omics. Cell Genomics 2, (2022).

99. Schep, A. N., Wu, B., Buenrostro, J. D. & Greenleaf, W. J. chromVAR: inferring transcription-factor-associated accessibility from single-cell epigenomic data. Nat. Methods 14, 975–978 (2017).

